# Culture-Associated Changes in the Human Cardiac Fibroblast Surfaceome: A Proteomic Resource for Cell Surface Marker and Target Discovery

**DOI:** 10.64898/2026.11.09.765497

**Authors:** Roneldine Mesidor, Linda Berg Luecke, Matthew Waas, Jennifer Schneider, Garam Park, Ella Rowe, Kyu Shik Kim, Heather C. Jensen-Smith, Erin N. Schuette, Pranav Renavikar, Claudius Mahr, Daniel R. Anderson, Rebekah L. Gundry

**Author notes:** Corresponding author: Rebekah L. Gundry, PhD, University of Nebraska Medical Center, Department of Cellular and Integrative Physiology, 985850 Nebraska Medical Center, Omaha, NE, 68198-5850, USA. These authors contributed equally (Roneldine Mesidor, Linda Berg Luecke).

## Abstract

Cardiac fibroblasts (CF) are central to cardiac homeostasis and fibrosis, but the lack of reliable cell surface markers limits their direct isolation, characterization, and selective targeting in human myocardium. Most CF studies use cultured cells, yet how the cell surface changes during culture remains poorly defined. We used mass spectrometry-based surfaceome profiling in five sequential substudies of human CF, spanning failing and non-failing hearts, fresh and cryopreserved myocardium, and laboratory-isolated and commercially sourced cells. Culture was associated with extensive changes in surface protein abundance, which were largest between P0 and P1 and were reproduced in CF from fresh and cryopreserved tissue. These changes included increases in proteins associated with fibroblast activation and matrix production, and changes in proteins used as CF markers and pharmacologic targets. Whole cell proteomics showed that these changes extended beyond the cell surface. Because culture history shapes cultured CF, we examined 30 years of CF literature, revealing widespread use of later, mixed, or unreported passages and incomplete reporting of culture conditions. To aid future studies, we developed CFxplorer (CFxplorer.net), an interactive resource that integrates this literature analysis, passage-resolved protein trajectories, and a Human CF Surfaceome Atlas of 1,577 cell surface N-glycoproteins detected across 234 CF samples, with extracellular peptide evidence and biological and pharmacologic annotations to help prioritize candidate surface markers and targets for validation. Collectively, this study expands the human CF surfaceome, defines how it changes during culture, and shows that the properties of cultured CF should be verified, not assumed, before use in experiments.

## 1. Introduction

Cardiac fibroblasts (CF) contribute to structural and electromechanical heart function and mediate intercellular communication with surrounding cells and the extracellular matrix (ECM) [1]. In response to cardiac injury, CF become activated and increase ECM protein production to maintain tissue integrity [1,2]. Pathological CF activation, common in heart failure, causes excessive ECM deposition, adverse ECM remodeling, and cardiac fibrosis [3–5], resulting in heart stiffness and impaired ventricular function [1,6]. Although pharmacological therapy can reduce overall mortality and improve symptoms [7], fibrosis remains largely irreversible [8], highlighting the need to better understand pathological fibrosis and develop targeted anti-fibrotic strategies [9,10].

Studying and selectively targeting human CF is limited by the lack of specific cell surface markers. Many commonly used CF markers are intracellular or ECM proteins [2,5], which limits their use for isolating viable cells, tracking CF in situ, and delivering imaging or therapeutic agents. No validated cell surface marker or marker combination currently enables sensitive and specific, non-genetic identification of homeostatic or activated primary human CF [5,11]. Systematic characterization of the human CF cell surface proteome (surfaceome) could identify candidates for antibody development, prospective isolation, culture assessment, and selective targeting, and define the receptors and adhesion molecules through which CF interact with their environment.

Without surface markers for direct isolation, most studies obtain CF by explantation and culture [12–16]. Tissue is dissociated, cells are plated under conditions that favor fibroblast outgrowth, and passaging expands cell numbers and helps separate CF from other cell types [11,17]. However, culture and serial passaging alter fibroblast phenotypes [18,19], including conversion to myofibroblasts, loss of *in vivo* ECM features [20–24], altered surface protein abundance [21,23], and diminished disease-associated differences [25]. These studies mostly examined selected proteins, often limited by antibody availability, thus the extent and timing of culture-associated changes across the cell surface remain unknown.

Here, we used CellSurfer [26], a mass spectrometry (MS)-based platform that identifies extracellularly exposed N-glycopeptides of cell surface N-glycoproteins, to assess the human CF surfaceome and how it changes in culture. In five sequential substudies, each designed in response to the one before, we asked when surface changes occur, whether CF from failing and non-failing hearts differ at the earliest passage, and how initial adhesion time, tissue cryopreservation, and commercial sourcing affect the surfaceome. We then examined how these culture variables have been used and reported across 30 years of CF literature. Finally, we integrated these data with a human CF Surfaceome Atlas into CFxplorer (CFxplorer.net), a public resource providing protein-level culture trajectories, extracellular peptide evidence, cardiac cell-type context, and available experimental and pharmacologic reagents. Our data show that the molecular features of cultured CF depend on culture history and should be characterized rather than assumed. To support this, CFxplorer allows investigators to interrogate these data and examine how proteins relevant to their own applications behave across culture conditions, providing a foundation for refining in vitro models and improving the translational relevance of CF research.

## 2. Materials and Methods

Reagent sources and catalog numbers are listed in Supplementary Table 1. Experimental details for flow cytometry, histology, and microscopy are provided in Supplementary Methods and Supplementary Table 2.

### 2.1. Patient donors used for CF isolation

Donors, tissue handling, culture conditions, and proteomic methods for each substudy are summarized in Figure 1. For Substudies I-IV, human cardiac tissue for CF isolation was obtained from institutional tissue banks at the Medical College of Wisconsin and University of Nebraska Medical Center under respective IRB approvals (PRO00025506; 0099-24-EP). Donors provided informed consent without compensation; de-identified clinical data are provided in Supplementary Table 3. Substudy V used cryopreserved CF from Lonza (CC-2904).

**Figure 1.**
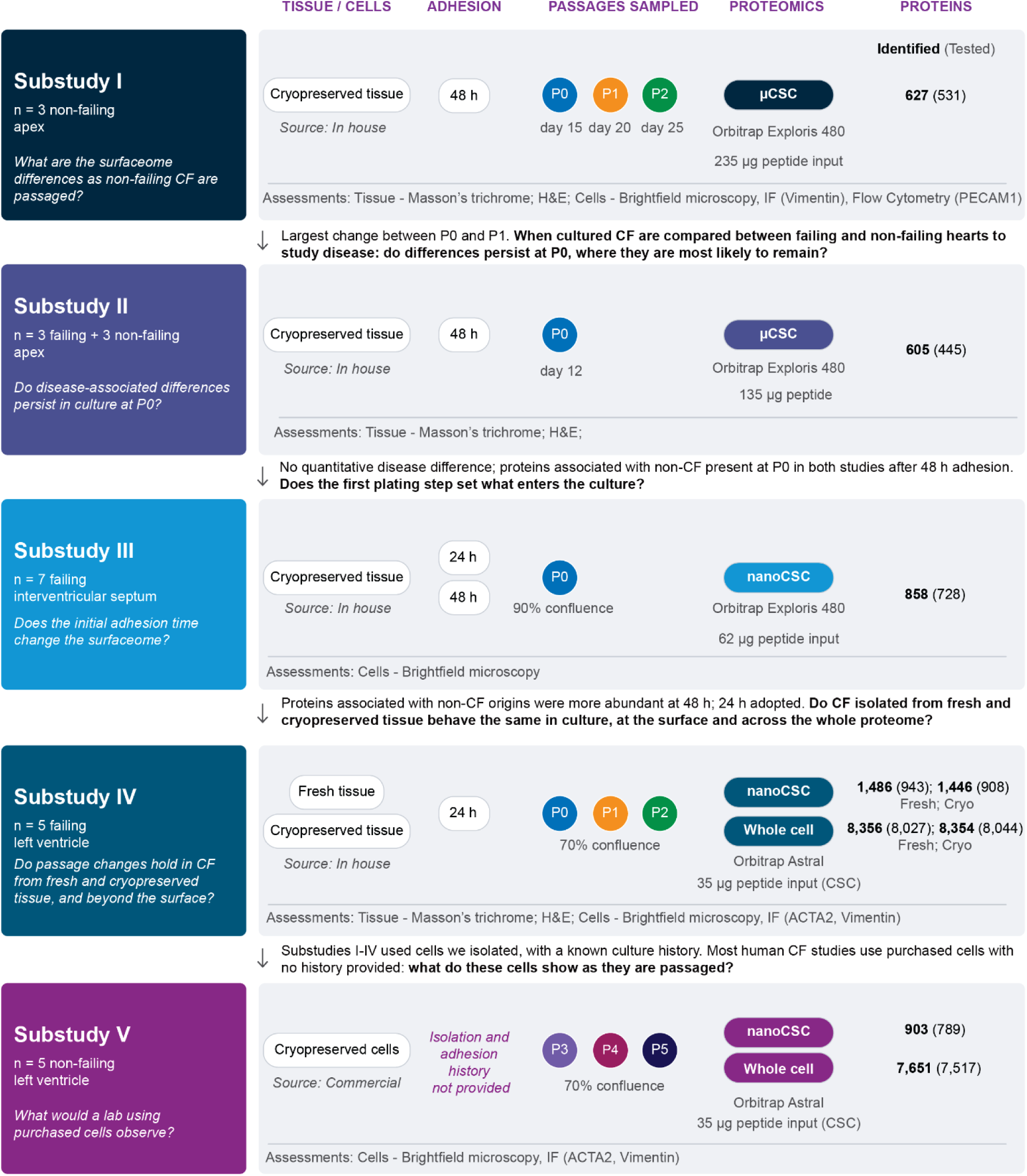
Design of the five substudies, each prompted by the results of the previous one. Each row summarizes one substudy: the donors and heart region, the question it addressed, the starting material and its source, the initial adhesion time before the first media change, the passages sampled, the proteomic methods, and the number of proteins measured. Source indicates whether CF were isolated in our laboratory or purchased. Protein counts are for cell surface N-glycoproteins classified as High or Medium confidence by Veneer (CSC) and for all proteins in whole cell lysates. Each count is the number identified at 1% FDR, with the number quantified in enough donors for statistical testing in parentheses. Assessments of tissues and cells are listed at the bottom of each row. Arrows between rows indicate the main finding of each substudy and the question it raised for the next. Abbreviations: IF, immunofluorescence; ACTA2, α-smooth muscle actin; PECAM1, platelet endothelial cell adhesion molecule 1; FDR, false discovery rate; P, passage; H&E, hematoxylin and eosin.

### 2.2. Isolation of primary CF from cryopreserved and fresh heart tissues

For Substudies I-IV, CF were isolated and cultured as described [27–29], with exceptions where noted. Upon receipt in the research laboratory, tissues for Substudies I-III were immediately cryopreserved in 10% DMSO, 10% FBS, and 80% DMEM until use. For Substudy IV, tissues were divided: one portion was processed for immunohistochemistry, one portion was used as fresh tissue, and one portion was cryopreserved until use. Tissues were thawed (if applicable) and cut into 1 mm^3^ pieces while submerged in Hanks Balanced Salt Solution with magnesium and calcium (HBSS+/+). Tissue pieces were placed in digestion solution (0.1 Units/mL collagenase B, 150 Units/mL collagenase IV, 0.5 mg/mL Dispase II, and 5 Units/mL DNase I in HBSS+/+) in a 50 mL conical tube at 37 °C while rotating end-over-end, then filtered with a 70 µm cell strainer. Flow through was collected and centrifuged at 300 x *g* for 5 minutes. Cells were resuspended in IMDM with 100 U/mL penicillin, 100 μg/mL streptomycin, 0.25 μg/mL amphotericin B, and 20% FBS and plated on 0.2% gelatin. For Substudies I and II, cells were plated for 48 h (i.e., adhesion time) before media was refreshed to remove non-adherent and dead cells and debris. For Substudy III, cells were plated for 24 or 48 h, and for Substudy IV for 24 h, before the first wash. For Substudy V, cells were cultured per manufacturer’s instructions to ensure findings would be representative of what other product users could expect. Brightfield imaging was used to evaluate cell morphology.

### 2.3. Cell Surface and Whole Cell Proteomics

CellSurfer combines cell surface capture [30], MS, and standardized bioinformatic curation to identify and quantify extracellular N-glycopeptides from surface glycoproteins [26]. Because methods and instruments improved during the study, each substudy used the most current workflow available: microscale cell surface capture (µCSC) [26] for Substudies I-II and a more sensitive version, nanoscale CSC (nanoCSC), for Substudies III-V (Supplementary Methods; Supplementary Figure 1), with an Orbitrap Exploris 480 MS for Substudies I-III and an Orbitrap Astral MS for Substudies IV-V. Whole cell proteomics for Substudies IV-V was performed as described [31] on an Orbitrap Astral MS (Supplementary Methods; Supplementary Table 4). Quantitative comparisons were made only within each substudy.

### 2.4. MS data analysis

Raw MS data were processed in Spectronaut™ 21 (Biognosys, Switzerland) using directDIA [32], and precursor-level reports were exported without normalization or imputation. Precursors passing precursor and protein Q-value ≤ 0.01 and posterior error probability ≤ 0.01 were used to calculate protein quantities with the MaxLFQ algorithm [33] in the iq R package [34]. Surfaceome data were restricted to High or Medium confidence cell surface proteins classified by Veneer [26,35]. Repeat injections were combined by their median, and one Substudy IV surface sample with a failed acquisition (donor 1376, fresh tissue, P0) was excluded.

Normalization was chosen for each dataset with an empirical diagnostic (Supplementary Methods). All datasets were median-normalized except Substudy III CSC, whose sample-wide shift followed adhesion time consistently across donors and would have been removed by normalization. The fresh and cryopreserved arms of Substudy IV were normalized and analyzed separately. Identified proteins passed a 1% FDR at the precursor and protein levels in at least one sample; quantified proteins had an abundance value in a given sample; and tested proteins were quantified in enough donors in every condition to enter the statistical model (Supplementary Methods). Missing values were not imputed; for proteins that were not tested, we report the number of donors in which each was quantified.

Differential abundance was tested separately for each dataset with limma [36,37], modeling log_2_ abundance on condition with donor as a blocking factor (Substudy II, which compared different donors, was unpaired) and robust, intensity-trended empirical Bayes moderation [38,39]. Each protein was tested across all conditions with the moderated F-test, and P values were adjusted by the Benjamini-Hochberg method [40]; FDR < 0.05 was considered significant and 0.05 ≤ FDR < 0.10 suggestive. DEqMS [41] was used as a sensitivity analysis. Descriptive analyses of timing, donor coordination, variance, trajectories and arm agreement are described in the Supplementary Methods. Analyses used R 4.6.1, limma 3.68.5, DEqMS 1.30.0 and e1071 1.7.17 [36,41,42]. Details are in the Supplementary Methods.

### 2.5. CFxplorer development and data integration

CFxplorer.net was developed with an iterative natural-language programming workflow as a React and TypeScript application built with Vite, with data services provided through an Express-based application programming interface. In addition to the literature and proteomic datasets, CFxplorer incorporates data, annotations, and links from the UniProt Knowledgebase, Human Protein Atlas [43], ABCD Antibody Database [44], Antibody Registry [45], Protter [46], Phobius [47], TMHMM [48], SOSUI [49], CellChatDB [50], CellPhoneDB [51], GPCRdb [52], ChEMBL [53], the Broad Institute Drug Repurposing Hub [54], DrugBank [55], PubMed, Gene Ontology [56,57], EMBL-EBI QuickGO [58], the Kyoto Encyclopedia of Genes and Genomes (KEGG) [59], and two published single-cell/single-nucleus RNA-seq (sc/snRNA-seq) analyses of the human heart [60,61].

### 2.6. Cell-type signatures, gene-set analyses, pathway scores and ligand-receptor analyses

Each analysis used one protein group per gene in each dataset. For within-donor change and signatures, tested groups were used first, then the group quantified in the most samples; for gene-set and ligand-receptor analyses, which use limma statistics, the tested group with the lowest omnibus P value; and for pathway scores, the group quantified in the most samples, then the one with the lowest omnibus P value. Change was the within-donor difference in log_2_ abundance between the first and last conditions (P0 and P2 in Substudies I and IV, P3 and P5 in Substudy V, 24 h and 48 h in Substudy III). For Substudy II, each failing donor was compared with the median of the non-failing donors, using proteins quantified in at least two non-failing donors. A protein quantified at only one endpoint was recorded as a gain or loss of detection.

Cell-type and fibroblast-state signatures were derived from the adult human heart atlas [62], obtained from CZ CELLxGENE Discover [63]. Raw counts were summed per donor and group (author cell-type annotation; unassigned cells and doublets excluded) when the group had at least 20 cells in that donor. Pseudobulk counts were converted to log_2_(counts per million + 1). Each cell type was compared with all other cell types, and each fibroblast state (FB1–FB7) with the other fibroblasts, by Welch’s t-test across donor pseudobulks, with Benjamini-Hochberg adjustment. Markers had a log_2_ fold change ≥ 0.5, adjusted P < 0.05, and at least 1 count per million in at least half of the group’s donors. Each signature comprised the 10 highest-ranked markers, by fold change, whose proteins were tested at the cell surface in Substudy I or IV. FB6 and FB7 were excluded because their top markers duplicated the endothelial and cardiomyocyte signatures. A signature’s score for a donor was the mean within-donor change of its members quantified at both endpoints, requiring at least three members.

For integrated gene-set analysis for each contrast, every tested protein (one per gene) was assigned z-score from its limma result, z = sign(log_2_ fold change) × Φ^−1^(1 - P/2). The fresh and cryopreserved arms of Substudy IV came from the same donors, so their scores were combined as (z_fresh_ + z_cryo_)/√(2 + 2r), where r is the correlation between the arms. Culture-associated change was defined from Substudies I and IV (P2 versus P0); Substudies II, III, and V were analyzed separately and compared with it. CellSurfer data were analyzed with mitch [64], a multi-contrast rank-based MANOVA of Substudies I and IV. Whole cell data, available for Substudy IV, and each individual contrast were analyzed with fgsea. Gene sets were MSigDB 2026.1 Hallmark [65], Reactome, GO biological process sets, and matrisome sets [66]. Sets had 5-300 (surface) or 10-500 (whole cell) tested members, with the tested proteins as background; P values were adjusted by the Benjamini-Hochberg method. Agreement between datasets was the percentage of gene sets with normalized enrichment scores (NES) of the same sign, for all sets and for sets with adjusted P < 0.25 in both.

Per-sample pathway scores were calculated with GSVA [67] (Gaussian kernel) on proteins quantified in every sample of a dataset, requiring at least 5 (surface) or 10 (whole cell) set members. CellSurfer and whole cell data, and the two arms of Substudy IV, were scored separately. Passage effects were tested within each dataset with a one-way ANOVA with donor as a blocking factor, and the timing of change with P1-versus-P0 and P2-versus-P1 contrasts from the same model; results were unchanged with donor as a random intercept.

Ligand-receptor network pairs were obtained from the Cell-Cell interaction database (v1.0) [68], keeping ligand or extracellular matrix–receptor pairs and receptor-receptor pairs. A surface protein was considered changed with culture when its combined Substudy I and IV analysis had FDR < 0.05, its direction was the same in Substudy I and both arms of Substudy IV, and its mean |log_2_ fold change| was ≥ 1. Pairs were drawn when both partners were tested at the surface and both had changed. The outer ring assigned proteins to pathways chosen from Reactome, matrisome, and Hallmark sets of ≤ 150 genes. Sets were added in turn, each the one covering the most proteins not yet assigned, until eight were chosen or no set added at least two. CFxplorer’s Ligand-Receptor page uses CellChatDB and CellPhoneDB (v5.0.0) pairs, scored within each sample from the abundance ranks of both partners. These analyses used R 4.3.3 [42] with limma 3.58.1 [36,37], mitch 1.23.1 [64], fgsea 1.39.4 [69], GSVA 1.50.0 [67] and circlize 0.4.15 [70], and Python 3.13 with anndata 0.13.4 [71] and SciPy 1.18.1 [72] for the atlas pseudobulk analysis.

### 2.7. Literature Search

We surveyed PubMed for primary research articles published between January 1, 1994 and June 11, 2024 that used isolated, cultured cardiac fibroblasts from human, mouse, or rat. From these, we manually extracted data for 13 experimental details related to CF isolation and culturing from 1,069 primary research articles that were in English and had not been retracted. Because details for “cell density at the time of experimentation” were reported using highly variable formats, the extracted values were assigned to standardized categories to enable quantitative summarization across studies. All details are available for review and download through CFxplorer.

## 3. Results

### 3.1. Study overview

We conducted five substudies; each designed in response to the findings of the previous substudy (Figure 1). Substudy I defined surfaceome trajectories across the first three passages (P0-P2) of CF from non-failing hearts. Because the largest changes occurred after P0, Substudy II compared CF from failing and non-failing hearts at P0, without passaging. Proteins commonly used as endothelial markers were readily detected at P0 in both substudies after a 48 h initial adhesion period, so Substudy III tested whether shortening this period to 24 h altered the surfaceome. Substudy IV adopted the shorter adhesion period, compared CF isolated from matched fresh and cryopreserved portions of failing myocardium, and added whole cell proteomics. Because most human CF studies use purchased cells (Section 3.8), Substudy V profiled commercially sourced CF across P3-P5.

Across substudies, cells showed the spindle-shaped morphology typical of fibroblasts, with no apparent differences among donors or passages (Supplementary Figure 2), and were vimentin (VIM)-positive in Substudies I, IV, and V; α-smooth muscle actin (ACTA2), assessed in Substudies IV and V, was present in a subset of cells (Supplementary Figure 3A-C). Histology showed minimal to mild interstitial fibrosis in the two Substudy I donors with available tissue, moderate interstitial fibrosis in one additional non-failing donor, and mild to moderate interstitial fibrosis in three failing donors from Substudy II, and minimal to moderate interstitial fibrosis in the failing left ventricular tissue of all five Substudy IV donors (Supplementary Figure 4). Missing values in the MS data occurred mainly at low abundance, consistent with values below the detection limit, and were not imputed (Supplementary Figure 5A-D).

### 3.2. Substudy I: The human CF surfaceome changes most between P0 and P1

We first compared CF from three non-failing hearts at P0, P1, and P2 (Figure 1, Substudy I). CellSurfer identified 627 cell surface N-glycoproteins, of which 531 were tested and 467 were quantified in every sample. Samples separated by passage along PC1 (38.6% of variance), with P0 apart from P1 and P2 in every donor (Figure 2A), and most proteins changed more between P0 and P1 than between P1 and P2 (60% of 526 proteins; median absolute change 0.42 vs 0.29 log₂; Figure 2B). Clustering of within-donor changes grouped these proteins into three trajectories: a steady decrease (Cluster 1: 196 proteins), a late increase between P1 and P2 (Cluster 2: 168) and an early increase between P0 and P1 (Cluster 3: 162) (Figure 2C). Fourteen proteins changed significantly with passage (FDR < 0.05). Thirteen decreased, including eight proteins commonly used as endothelial markers (VWF, PECAM1, CALCRL, FLT1, MMRN2, PLVAP, CD34, and ICAM2) together with ACAN, AOC3, TSPAN15, COL6A1, and COL6A2; PECAM1 and VWF fell more than 15-fold in every donor (Figure 2D). The one significant increase was HHIP (+3.2 log₂), and other surface proteins, including LGR4, HAPLN1, and VIPR1, rose by a median of more than 2 log₂ without reaching significance with three donors (Figure 2E). Proteins commonly used to identify CF also rose modestly, including THY1 in all three donors and FAP and PDGFRA in two of three. Consistent with the MS data, flow cytometry identified a small PECAM1-positive population that decreased with passage (22% to 4%; Supplementary Figure 6).

**Figure 2.**
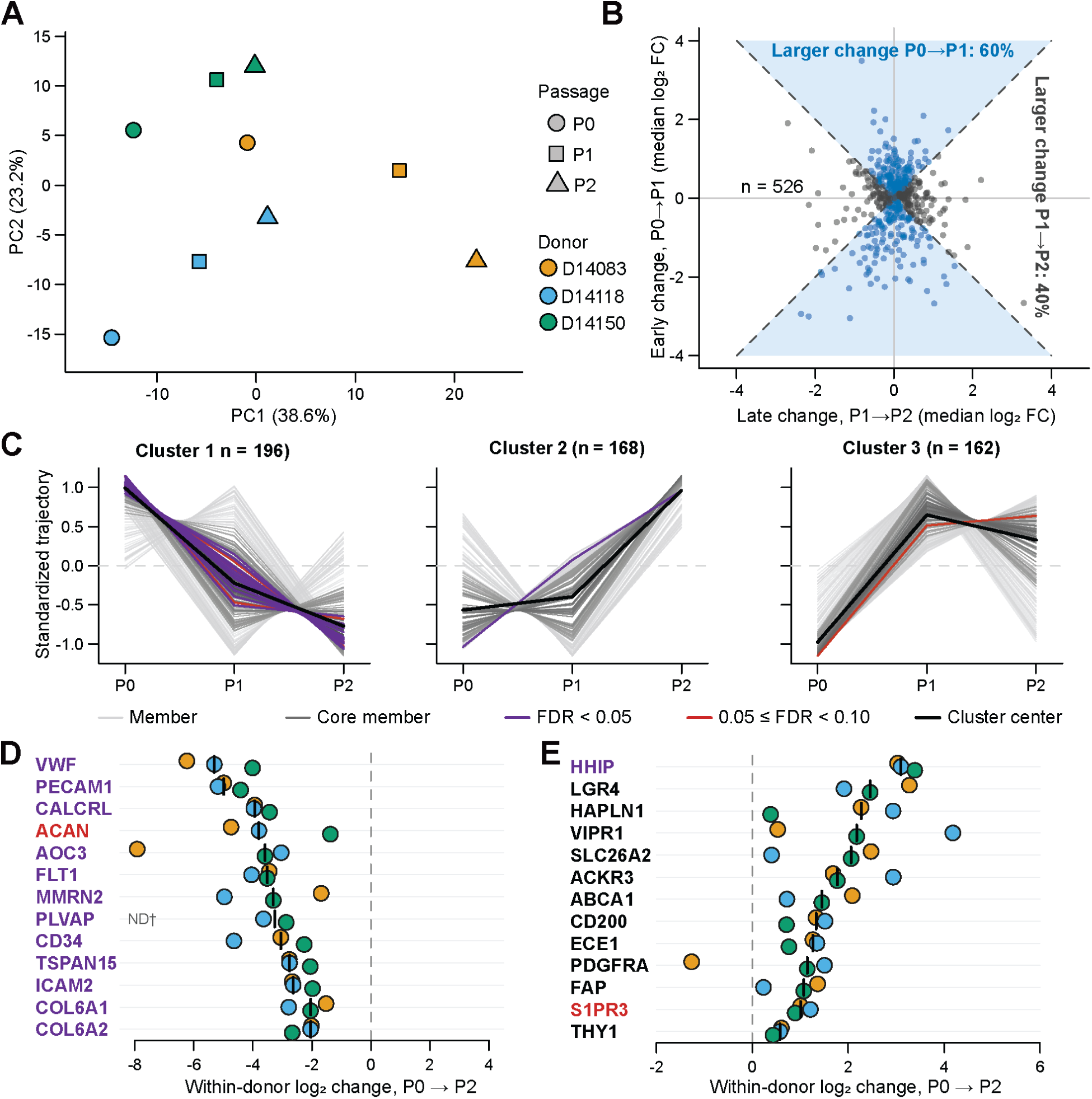
Surfaceome of human CF across P0–P2 (Substudy I). A. Principal component analysis of cell surface protein abundance in CF from three non-failing hearts at P0, P1, and P2, using proteins quantified in every sample. Color indicates passage and symbol indicates donor; lines connect samples from the same donor. B. Timing of change. For each protein with both intervals measured in at least two donors, the median within-donor log₂ change from P0 to P1 (early, y-axis) is plotted against that from P1 to P2 (late, x-axis). Dashed lines mark equal magnitudes of early and late change; shading and blue points mark proteins with a larger change from P0 to P1 than from P1 to P2. C. Trajectory clusters from fuzzy c-means clustering (k = 3) of within-donor log₂ changes from P0. Each line is one protein’s median trajectory, standardized to mean 0 and SD 1; light gray, cluster members; dark gray, core members (membership ≥ 0.5); green, omnibus FDR < 0.05; orange, 0.05 ≤ FDR < 0.10; black, cluster center. D-E. Within-donor log₂ change from P0 to P2 for proteins that decreased with omnibus FDR < 0.05 (D) and for selected proteins that increased (E). Each point is one donor, and the black bar marks the median; gene names in green have FDR < 0.05. ND†, loss of detection in too few donors to mark.

### 3.3. Substudy II: CF from failing and non-failing hearts at P0

Because the largest changes in Substudy I occurred after P0, we compared CF from failing and non-failing hearts at P0, the earliest available point (Figure 1, Substudy II). CellSurfer identified 605 cell surface N-glycoproteins, and all 445 tested proteins were quantified in all six samples. No protein differed significantly between failing and non-failing CF after correction for multiple testing (limma, Benjamini–Hochberg FDR < 0.05). Before this correction, 39 proteins had a nominal P < 0.05 (Supplementary Figure 7A), close to the 22 expected by chance among 445 tests (Supplementary Figure 8A). The groups differed more clearly in which proteins were detected. Twenty proteins were detected exclusively in CF from failing hearts, 13 of which in at least two of three failing donors, including TNFAIP6, SGCA, and SLC8B1 in all three, and SFRP4, VCAM1, PODN, and THBS3 in two donors (Supplementary Figure 7B). Therefore, the surfaceome of CF from failing hearts differed from that of non-failing CF, but more modestly than expected. With three donors per group, and similar variation among donors in each group (median per-protein SD 0.58 and 0.50 log₂ in failing and non-failing CF, respectively), this design could detect only large differences in abundance. Limited sensitivity alone seems unlikely to explain this result because the same µCSC workflow detected disease-associated differences in the surfaceome of acutely isolated (i.e., not plated or cultured) cardiomyocytes from three failing and three non-failing hearts [26]. Differences between failing and non-failing CF have also been reported to diminish in culture in canine CF [25], suggesting that the steps before P0 may already influence the CF surfaceome. One such step is the initial adhesion period: in Substudies I and II, cells were plated for 48 h before the first wash, and proteins associated with non-CF cell types, including proteins commonly used as endothelial markers, were detected in every donor at P0. Therefore, we then investigated how the length of the initial adhesion period affects the surfaceome.

### 3.4. Substudy III: A longer initial adhesion period increases proteins commonly used as endothelial markers

Substudy III compared CF from seven failing donors washed after 24 h or 48 h, the time before non-adherent cells are first removed, and examined both the surfaceome as a whole and the proteins commonly used as endothelial markers (Figure 1, Substudy III). CellSurfer identified 858 cell surface N-glycoproteins, of which 728 were tested. Surface protein abundance was higher overall after 48 h in six of seven donors (median increase in sample median, 0.41 log₂; Supplementary Figure 7C), so these data were not normalized. Fifty-eight proteins were more abundant after 48 h (FDR < 0.05), and none were less abundant (Supplementary Figure 7D). The largest increases exceeded the sample-wide shift, up to 5-fold for MFAP5, and included VTN and BMP1 (Supplementary Figure 7D, E). The endothelial marker proteins CD34 (+1.8 log₂), VWF (+1.5) and ANGPT2 (+0.8) were more abundant after 48 h, as were AOC3 (+1.0) and CD163 (+2.0), a protein associated with non-CF cell types (FDR < 0.05; Supplementary Figure 7D, E). The response was not uniform: PECAM1 rose only slightly (+0.6 log₂; FDR = 0.08; Supplementary Figure 7E), and CDH5, ENG, and CD14 did not change significantly. Nine proteins were detected only at 48 h, but only GPM6B in a majority of donors (5 of 7; Supplementary Figure 7F). Therefore, a longer initial adhesion increased several proteins commonly used as endothelial markers, although individual proteins responded differently and none can be read as a direct measure of cell composition. Because CF cultures were established reliably after 24 h, we used the shorter period in Substudy IV.

### 3.5. Substudy IV: CF from fresh and cryopreserved myocardium follow shared culture trajectories

Human myocardium for research becomes available unpredictably, so tissue is often cryopreserved as it arrives and processed later in batches. To test whether this affects culture-associated change, Substudy IV divided failing myocardium from five donors at collection, using one portion immediately for CF isolation and cryopreserving the other before isolation (Figure 1, Substudy IV). Cells were plated for 24 h before the first wash and analyzed at P0, P1 and P2.

CellSurfer identified 1,486 and 1,446 N-glycoproteins in the fresh and cryopreserved arms, of which 943 and 908 were tested, and 554 and 403 changed with passage (FDR < 0.05). In both arms, P0 samples separated from P1 and P2 along PC1 (Figure 3A; Supplementary Figure 8C), and most proteins changed more between P0 and P1 than between P1 and P2 (75% fresh, 68% cryopreserved; Figure 3B). Individual proteins illustrate this: SEMA7A and THY1 increased and CD36 and AOC3 decreased at the cell surface in both arms (Figure 3D; whole cell data in Supplementary Figure 9A). The agreement held across the surfaceome: fold changes from P0 to P2 were strongly correlated across the 878 proteins tested in both arms (ρ = 0.82), and of the 401 of these that changed in the cryopreserved arm, 96% changed in the same direction in the fresh arm (Figure 3C; Supplementary Figures 10B and 11A-D). The arms were not identical, however: SEMA7A increased more, changes were more closely coordinated among donors (Section 3.6), and more proteins were detected mainly at P0 in the fresh arm than in the cryopreserved arm (Supplementary Figure 5E). CF from fresh and cryopreserved tissue therefore followed similar trajectories but differed in the magnitude and coordination of change, so tissue handling should be reported when comparing studies.

**Figure 3.**
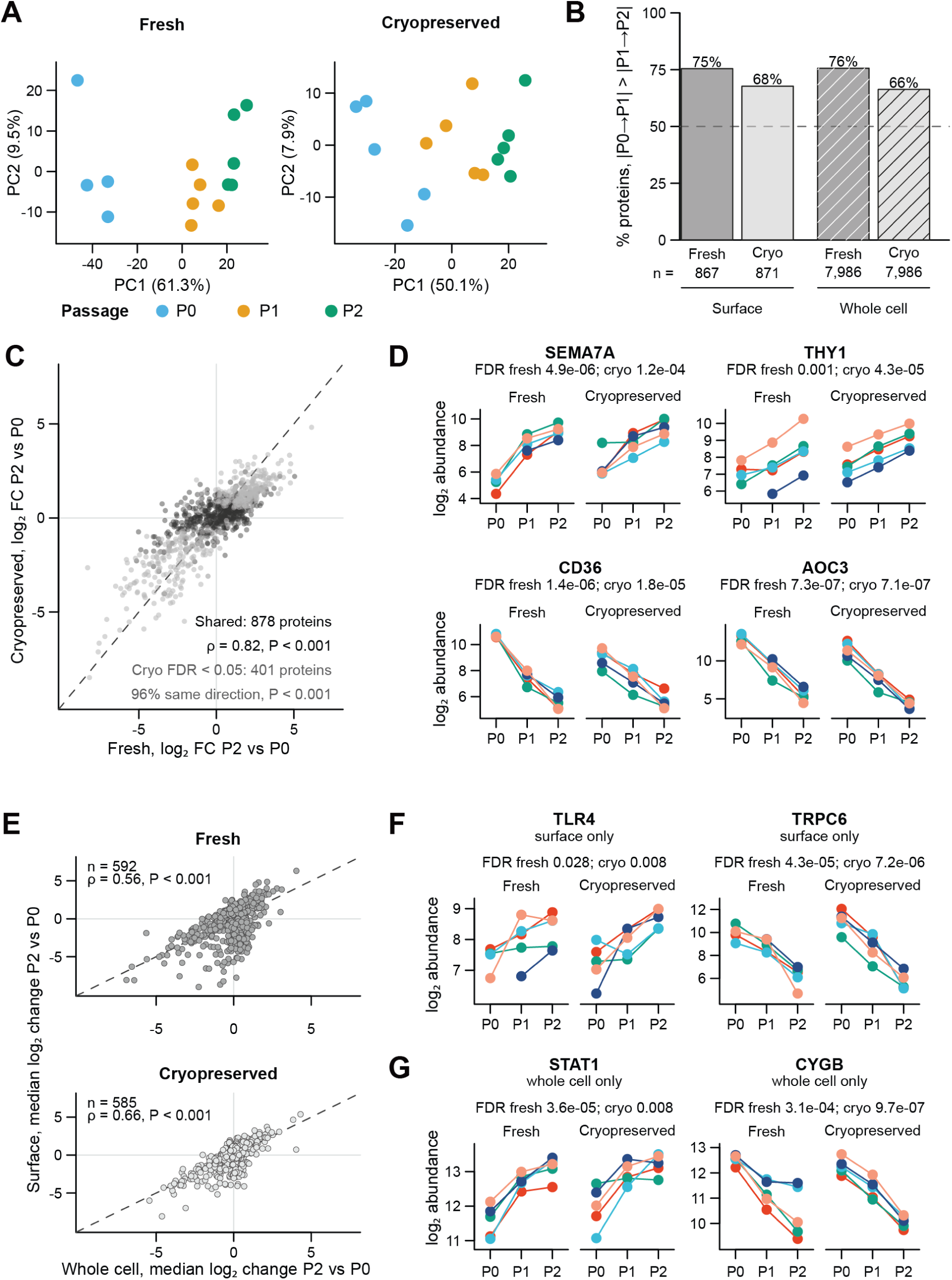
Surfaceome and whole cell proteome of CF isolated from matched fresh and cryopreserved myocardium across P0–P2 (Substudy IV). A. Principal component analysis of cell surface protein abundance, shown separately for the fresh and cryopreserved arms, using proteins quantified in every sample of that arm. Color indicates passage and symbol indicates donor; lines connect samples from the same donor. B. Percentage of proteins with a larger median within-donor change from P0 to P1 than from P1 to P2, for each arm and assay; the dashed line marks 50%. C. Log₂ fold change from P0 to P2 in the fresh and cryopreserved arms for surface proteins tested in both; blue, proteins with FDR < 0.05 in the cryopreserved arm; dashed line, identity. D. Cell surface log₂ abundance of SEMA7A, THY1, CD36, and AOC3 in each donor of the fresh (blue) and cryopreserved (orange) arms, with the omnibus FDR in each arm. E. Median within-donor log₂ change from P0 to P2 in surface and whole cell data for proteins evaluable by both approaches, in the fresh and cryopreserved arms; dashed line, identity. F-G Log₂ abundance in each donor for proteins quantified by only one approach, TLR4 and TRPC6 at the cell surface (F) and STAT1 and CYGB in whole cell data (G), shown as in D.

Whole cell proteomics extended these observations beyond the cell surface. Of 8,027 (fresh) and 8,044 (cryopreserved) proteins tested, 3,681 and 2,170 changed with passage, again mostly between P0 and P1 (76% and 66%, respectively; Figure 3B), and of the 2,152 that changed in the cryopreserved arm and were tested in both, 97% changed in the same direction in the fresh arm (Supplementary Figure 11E-H). Similar to the surface dataset, samples separated by passage along PC1 (59.2% of variance), with P0 distinct from P1 and P2 in every donor for fresh and cryopreserved samples (Supplementary Figure 8B, D). CellSurfer enriches cell surface N-glycoproteins and provides direct evidence of extracellular localization, whereas whole cell proteomics measures a broader, largely intracellular set of proteins. For proteins measured by both, changes from P0 to P2 were positively correlated (ρ = 0.56 fresh and 0.66 cryopreserved; Figure 3E; Supplementary Figure 10C,D), so surface and total abundance often changed in the same direction, but whole cell abundance did not reliably predict surface abundance. Each approach also captured proteins that the other did not. Among proteins tested only at the cell surface, TLR4 increased with passage and TRPC6 decreased (Figure 3F), whereas among those tested only in whole cell data, the intracellular proteins STAT1 and CYGB increased and decreased, respectively (Figure 3G). Whole cell data also showed whether cells acquired a contractile myofibroblast profile. ACTA2 decreased in every donor of the fresh arm (-1.3 log₂, FDR < 0.001) and in four of five donors of the cryopreserved arm (not significant), CNN1 did not change consistently, and TAGLN and MYL9 increased only modestly (≤ 0.8 log₂), consistent with detection of ACTA2 in only a subset of cells by imaging (Supplementary Figure 3B,C). Like Substudy I, these experiments used CF isolated in our laboratory with a known culture history. Most human CF studies instead use purchased cells, which may arrive at later passages, and without details on how they were isolated or cultured; therefore, we then investigated how commercially sourced CF change across passages.

### 3.6. Substudy V: In commercially sourced CF, variation among donors exceeds the effect of passage

Substudy V profiled commercially sourced CF from five non-failing donors at P3-P5, cultured according to the supplier’s instructions. P3 was the earliest passage that could be studied because cells were supplied at P2 (Figure 1, Substudy V). CellSurfer identified 903 cell surface N-glycoproteins (789 tested), and whole cell proteomics identified 7,651 proteins (7,517 tested). Changes with passage were small at the cohort level: only ISLR changed significantly at the surface (−1.7 log₂, FDR = 0.02), no whole cell protein reached FDR < 0.05, and passage explained 1.5% of the sample-wide shift in surface abundance and 11% in whole cell abundance. Variation was instead dominated by donor: passage explained more variance than donor for only 7% of surface and 8% of whole cell proteins, compared with 57-74% of surface proteins in the two arms of Substudy IV (Figure 4A; Supplementary Figure 9B, C). Donors also did not change in the same way. The changes in one donor from P3 to P5 were essentially unrelated to those of another (median pairwise ρ = 0.05 for surface and 0.06 for whole cell proteins, compared with 0.60–0.78 for surface proteins in Substudy IV; Figure 4B), and 37% of surface proteins changed in the same direction in at least four of five donors, no more than expected by chance (37.5%), compared with 51% of whole cell proteins (Supplementary Figure 10E).

**Figure 4.**
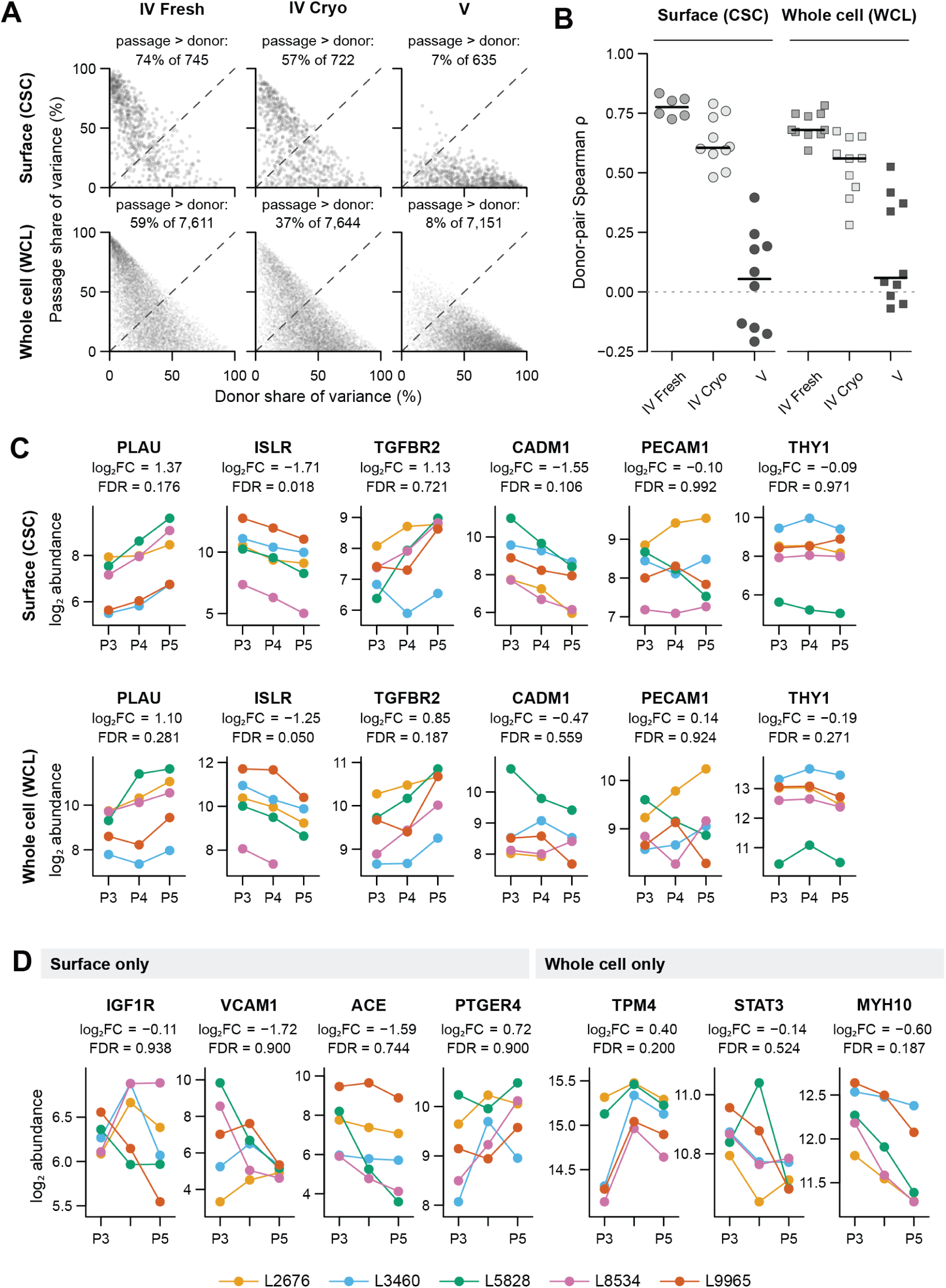
Surfaceome and whole cell proteome of commercially sourced CF across P3–P5 (Substudy V). A. Donor and passage shares of variance for each protein quantified in every sample, from a linear model containing both terms, for each arm of Substudy IV (P0–P2) and for Substudy V (P3–P5); cell surface (top) and whole cell (bottom) data; dashed line, equal shares. Each dataset was analyzed separately. B. Donor coordination. Each point is the Spearman correlation between the within-donor log₂ changes, from the first to the last passage, of two donors, using proteins quantified at both passages in every donor of every dataset of that assay; black bars mark medians. C. Log₂ abundance at P3, P4, and P5 for PLAU, ISLR, TGFBR2, CADM1, PECAM1, and THY1 at the cell surface (top) and in whole cell data (bottom), with the limma log₂ fold change (P5/P3) and omnibus FDR. Each line is one donor. D. Log₂ abundance at P3, P4 and P5 for proteins quantified by only one approach, IGF1R, VCAM1, ACE, and PTGER4 at the cell surface and TPM4, STAT3, and MYH10 in whole cell data, shown as in C.

Against this background, a few proteins changed consistently within donors (Figure 4C). ISLR decreased in every donor at the surface and in every donor with both passages measured in whole cell data (-1.2 log₂, FDR = 0.05), and PLAU increased in every donor at the surface and in four of five in whole cell data (+1.4 and +1.1 log₂) without reaching significance. CADM1 decreased at the surface in all five donors (-1.6 log₂) but changed little in whole cell data (-0.5). In contrast, THY1 remained stable, as did PECAM1, which was quantified at every passage alongside the endothelial marker proteins CD34, CDH5, and ENG; their presence is therefore not unique to CF isolated in our laboratory. Among proteins measured by only one approach, ACE decreased at the surface in every donor and MYH10 in whole cell data in four of five, but neither reached significance (Figure 4D). CNN1, which did not change consistently in Substudy IV, decreased in every donor in whole cell data (−0.8 log₂, FDR = 0.20). In commercially sourced CF at P3-P5, passage-associated change was therefore small and not shared across donors, although individual proteins could change consistently within donors. Because the supplier provided no isolation or culture history and commercial source and later passage cannot be separated in this design, these data do not establish why these cells vary so much among donors. They do show that studies using purchased cells should treat donor, or lot, as a major source of variation and should not assume that these cultures are free of proteins associated with non-CF cell types.

### 3.7. Across substudies, culture shifts CF away from in vivo cell-type and fibroblast-state markers and alters pharmacologic targets

Having characterized each substudy on its own, we next asked which changes were shared across substudies. We followed markers of cardiac cell types and fibroblast states, and pharmacologic targets, at the donor level (Figure 5), and combined substudy statistics in gene-set and ligand–receptor analyses (Figure 6). Because methods differed among substudies, these comparisons concern the direction and consistency of change, not abundance.

**Figure 5.**
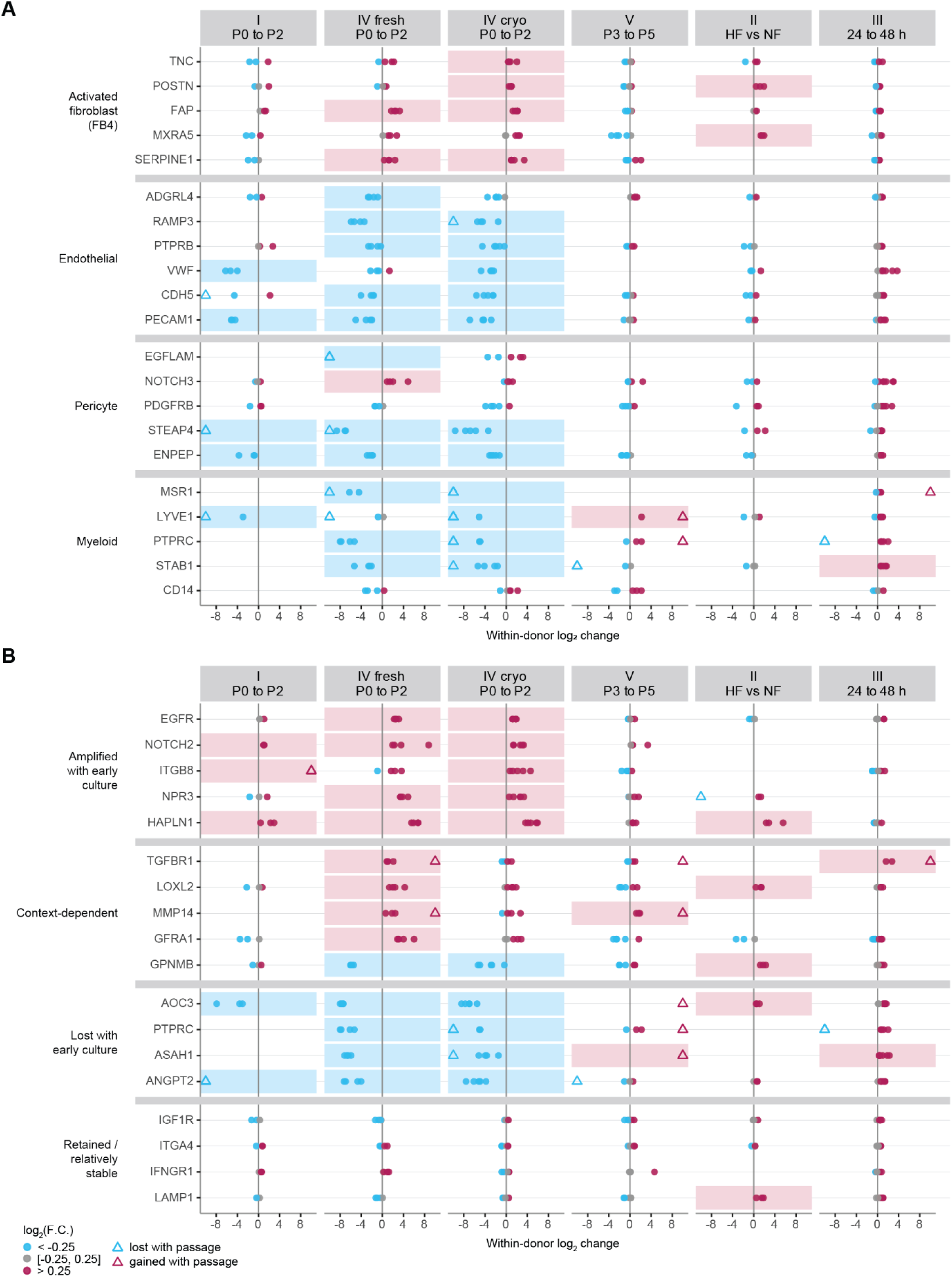
Donor-level changes in cell-type and fibroblast-state markers and pharmacologic targets at the cell surface. Each point is one donor. Columns show the within-donor change in log_2_ abundance from the first to the last passage (P0 to P2 in Substudies I and IV; P3 to P5 in Substudy V). Substudy II shows each failing donor relative to the median of the non-failing (NF) donors, and Substudy III the within-donor change from 24 h to 48 h initial adhesion. Point color indicates an increase (> 0.25 log_2_), a decrease (< -0.25 log_2_), or minimal change; shading marks proteins that changed by more than 0.25 log_2_ in the same direction in every donor, counting gain or loss of detection as an increase or decrease. Open triangles at the panel edge mark donors in which the protein was quantified at only one endpoint (gain or loss of detection). A. The five highest-ranked surface-tested markers of activated fibroblasts (FB4, versus other fibroblasts) and of endothelial cells, pericytes, and myeloid cells (versus other cell types) in the adult human heart atlas. PECAM1, a common marker of endothelial cells, is shown for reference (rank 15). All signatures are scored in Supplementary Figure 12. B. Surface proteins with approved or investigational drugs, grouped by pattern of change. Because methods differed among substudies (Figure 1), columns show the direction and consistency of change among donors, not abundance.

**Figure 6.**
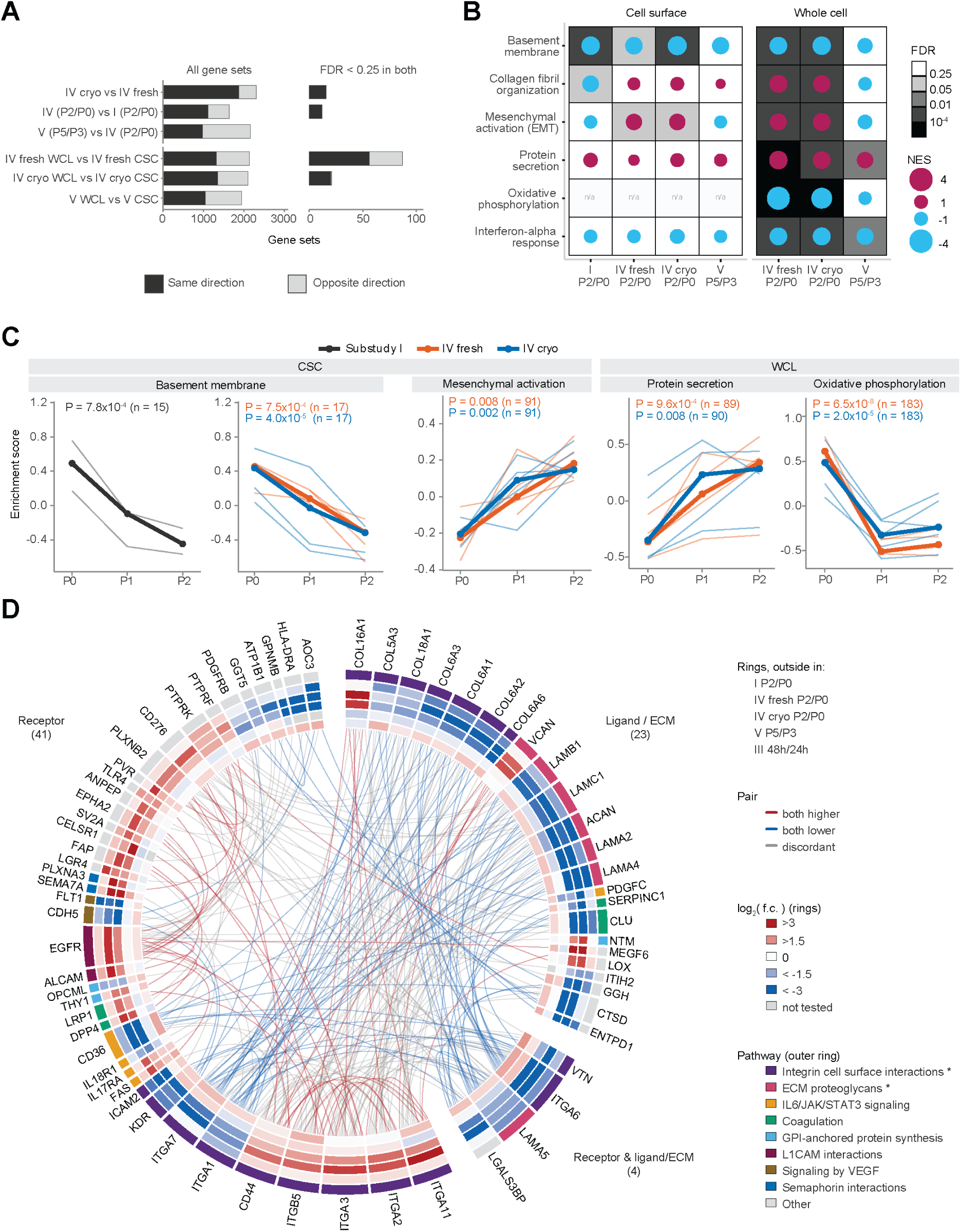
Agreement between datasets, integrated gene set enrichment, pathway scores, and ligand-receptor pairs across substudies. A. Number of gene sets with normalized enrichment scores (NES) of the same (dark) or opposite (light) sign in two datasets, between substudies at the cell surface (top) and between cell surface and whole cell data within a dataset (bottom), for all gene sets scored in both datasets (left) and for sets with FDR < 0.25 in both (right); n/a, no set passed. The fresh and cryopreserved arms of Substudy IV were combined at the protein level before enrichment. B. Gene set enrichment (fgsea) of each contrast for six highlighted pathways at the cell surface (left) and in whole cell data (right). Tile shading shows the FDR, and dot size and color the NES (maroon, higher and blue, lower in the later passage); n/a, too few members tested. Pathways were chosen to represent the programs that changed with culture at the cell surface (Substudies I and IV combined) or in whole cell data (Substudy IV); all results are in Supplementary Table 6. C. Per-sample GSVA scores for basement membrane and mesenchymal activation (EMT) at the cell surface (CSC) and for protein secretion and oxidative phosphorylation in whole cell data (WCL). Thin lines, individual donors; thick lines, median of donors; grey, Substudy I; orange, fresh arm; blue, cryopreserved arm of Substudy IV. P values are from a one-way ANOVA with donor as a blocking factor within each dataset; n, proteins mapped to the pathway and quantified in every sample. Scores are compared within a dataset only. D. Ligand-receptor pairs from the Cell-Cell interaction database in which each partner changed by at least 2-fold, consistently in Substudies I and IV (meta-analysis FDR < 0.05). Outer ring, pathway assignment (* gene sets significant in the integrated surface analysis); inner rings, log_2_ fold change in Substudy I, IV fresh, IV cryopreserved (P2/P0), V (P5/P3) and III (48 h/24 h); grey, not tested. Links are red when both partners increased, blue when both decreased and grey when they changed in opposite directions. Pairs describe which partners are present at the surface and do not measure signaling.

To compare cultured CF with cells in the heart, we defined marker signatures from the adult human heart atlas [62]. For each of 11 cell types and 5 fibroblast states, marker genes were ranked by fold change, and the 10 highest-ranked genes whose proteins were tested at the cell surface in Substudy I or IV were retained (Supplementary Table 5).

Early culture shifted CF away from markers of other cardiac cell types and toward markers of an activated fibroblast state (Figure 5A). Among the top markers of activated fibroblasts (cluster FB4; a TGF-β-responsive fibroblast state in the adult heart atlas [62] whose markers (TNC, POSTN, FAP, SERPINE1, FN1, CCN2) overlap those of activated fibroblasts), FAP and SERPINE1 increased from P0 to P2 in every donor in both arms of Substudy IV, and TNC and POSTN increased in every donor in the cryopreserved arm. POSTN and MXRA5 were also higher in all three failing donors than in non-failing CF at P0, but none of these markers increased consistently across P3-P5 in commercially sourced cells. In contrast, the common endothelial markers ADGRL4, RAMP3, PTPRB, CDH5, VWF, and PECAM1, decreased in every donor, or all but one, in both arms of Substudy IV. Pericyte markers (ENPEP and STEAP4) and myeloid markers (PTPRC, STAB1, and MSR1) also decreased, several to below the detection limit, and PECAM1, VWF and ENPEP decreased in all three donors of Substudy I. The pericyte marker NOTCH3 was an exception, as it increased in every fresh-arm donor with a P0 sample.

Signature scores showed the same pattern (Supplementary Figure 12). From P0 to P2 in Substudy IV, the endothelial, pericyte, myeloid, and both cardiomyocyte signatures decreased in every donor, or all but one, of both arms. The activated fibroblast (FB4) signature increased in every cryopreserved-arm donor and in three of four fresh-arm donors and was higher in all three failing donors at P0. However, the fibroblast signature decreased in every fresh-arm donor and in four of five cryopreserved-arm donors (median -1.1 and -1.2 log_2_). Its members DCN, ABCA6, and CDH19 decreased in most donors, although NEGR1 increased in every donor and PDGFRA in every fresh-arm donor; as cultured CF acquired an activated profile they lost some, but not all, markers of fibroblasts in the heart. No signature changed in the same direction in every donor across P3-P5. In Substudy III, a longer initial adhesion increased most signatures, consistent with the sample-wide increase in that unnormalized dataset (median 0.41 log_2_; Section 3.4). Signature scores describe shifts in the measured surface profile, not cell composition.

Surface proteins targeted by approved or investigational drugs were also lost or gained during early culture (Figure 5B). AOC3 decreased in every donor of Substudy I (median −3.6 log_2_) and in both arms of Substudy IV (−7.6 and −7.0), as did ANGPT2, ASAH1, PTPRC, and GPNMB in Substudy IV. Conversely, EGFR, NOTCH2, NPR3, and HAPLN1 increased in every donor of both arms of Substudy IV, by up to median +6.3 log_2_ for HAPLN1. NOTCH2 and HAPLN1 also increased in all three donors of Substudy I and in four of five commercially sourced donors, and HAPLN1 and GPNMB were higher in all three failing donors at P0. AOC3, among the most abundant surface proteins at P0 in Substudies I and IV, was quantified in only one commercially sourced donor. Therefore, a target present on freshly isolated CF may be lost from cultured or purchased cells, while another becomes more prominent with passage.

To identify programs that changed consistently, we combined the passage responses of Substudies I and IV and compared them with Substudies II, III, and V. At the protein level, fold changes from P0 to P2 agreed between Substudies I and IV (ρ = 0.36 - 0.41 for surface proteins tested in both) and agreed more closely between the two arms of Substudy IV (ρ = 0.82) (Supplementary Figure 10A,B). They did not agree with changes across P3-P5 in commercially sourced cells (ρ ≤ 0.07) and were inversely related to the change after a longer initial adhesion (ρ = −0.32 to −0.47). Of 507 surface proteins tested in both Substudies I and IV, 279 changed with culture in a combined analysis (FDR < 0.05), and 241 of these changed in the same direction in Substudy I and both arms of Substudy IV.

Agreement at the pathway level mirrored that at the protein level (Figure 6A, Supplementary Figure 13). Gene sets changed in the same direction in 68% of the comparisons between Substudies I and IV, including all 12 sets with adjusted P < 0.25 in both. Agreement was 81% between the two arms of Substudy IV and 45% between Substudies IV and V, close to the 50% expected by chance. Surface and whole cell enrichment agreed for 62–64% of sets in Substudy IV. In the fresh arm, 36% of the 87 sets with adjusted P < 0.25 in both assays changed in opposite directions; therefore, whole cell changes did not reliably predict surface changes at the pathway level.

Gene-set analysis of the surface data identified 50 sets that changed with culture (FDR < 0.05), 38 which were in the same direction in Substudies I and IV (Figure 6B; Supplementary Table 6). Thirty of these 38 decreased, including basement membrane (FDR = 6 × 10^−4^), type VI collagen and laminin-containing sets, PDGF signaling, and platelet-associated sets. Those that increased included transporter, cell projection, and plasma membrane localization sets, driven by proteins such as EGFR, EPHA2, and ITGB1. Whole cell data from Substudy IV broadened this picture. Oxidative phosphorylation decreased sharply (NES -3.6, adjusted P = 6 × 10^−38^), as did interferon-α response (NES -2.2) and basement membrane (NES −2.2), whereas protein secretion (NES +2.1), collagen fibril organization (+2.2) and epithelial-mesenchymal transition (EMT; +2.0) increased (all adjusted P ≤ 0.002). Sets for canonical TGF-β signaling (Hallmark, Reactome and GO; adjusted P ≥ 0.81) did not change. Early culture was therefore associated with a secretory, matrix-producing profile and loss of oxidative metabolism rather than a canonical TGF-β response, consistent with the absence of a coordinated increase in ACTA2 (Section 3.5). Several of the programs that changed with passage were also higher in failing than in non-failing CF at P0 (basement membrane NES +2.2, collagen fibril organization +2.3, EMT +2.3; adjusted P ≤ 0.009). Across P3-P5, only whole cell protein secretion (+1.7) and interferon-α response (-1.9) changed in the same direction as in Substudy IV (adjusted P = 0.02).

Per-sample GSVA scores showed when these changes occurred (Figure 6C). In Substudy IV, scores changed more between P0 and P1 than between P1 and P2 in 15 of 22 dataset–pathway comparisons. Basement membrane was the main exception: its surface scores decreased in Substudy I (P = 7.8 × 10⁻⁴) and in both arms of Substudy IV (P = 7.5 × 10⁻⁴ and 4.0 × 10⁻⁵) and continued to fall between P1 and P2. Mesenchymal activation (EMT) scores increased at the surface in both arms (P = 0.008 and 0.002). In whole cell data, oxidative phosphorylation decreased (P = 6.5 × 10⁻⁸ fresh and 2.0 × 10⁻⁵ cryopreserved), and protein secretion increased (P = 9.6 × 10⁻⁴ and 0.008).

Finally, we placed the consistently changed surface proteins in a ligand–receptor network [68] (Figure 6D). Of 181 proteins that changed by at least 2-fold in the same direction in Substudies I and IV (105 increased, 76 decreased), 68 formed 245 ligand-receptor pairs in which both partners changed. Laminins and type VI collagens decreased together with their receptors ITGA6, ITGA7, and ITGA1, giving 72 pairs in which both partners were lower. Other receptors increased (ITGA11, ITGA2, ITGA3, ITGB5, CD44, EGFR), so 128 pairs had partners changing in opposite directions. Of these 68 selected proteins, 44 changed in the same direction across P3-P5, and 48 changed in the opposite direction after a longer initial adhesion. Pair membership describes which partners are present at the surface and does not measure signaling. These analyses can be repeated in CFxplorer, where the Signature Trajectory and Ligand-Receptor pages score any gene set or pair within a substudy, arm, and assay.

### 3.8. CF culture methods are heterogeneous and incompletely reported across 30 years of published studies

Because passage, initial adhesion time, and cell source each affected the CF surfaceome in our substudies, we asked how published work handled these variables. We extracted 13 experimental variables from 1,195 workflows reported among 1,069 primary research articles published over 30 years that used cultured human, mouse, or rat CF (Figure 7A). CF models and culture conditions varied considerably (Figure 7B). Passage, which produced the largest changes in our data between P0 and P1, ranged from P0 to P39: 23% of workflows used a single defined passage, 38% used cells beyond P2, 43% used a range or mixture of passages, and 35% did not report passage. Rodent studies mostly used acutely isolated cells, whereas human studies mostly used previously isolated cells, including commercially sourced or biobanked CF (Supplementary Figure 14A). Initial adhesion time, which affected proteins commonly used as endothelial markers in Substudy III, also varied, as did culture substrate and coating, basal medium, duration of culture, and cell density (Figure 7B). Several variables that affected the surfaceome here differ widely among published studies, and passage is often not reported.

**Figure 7.**
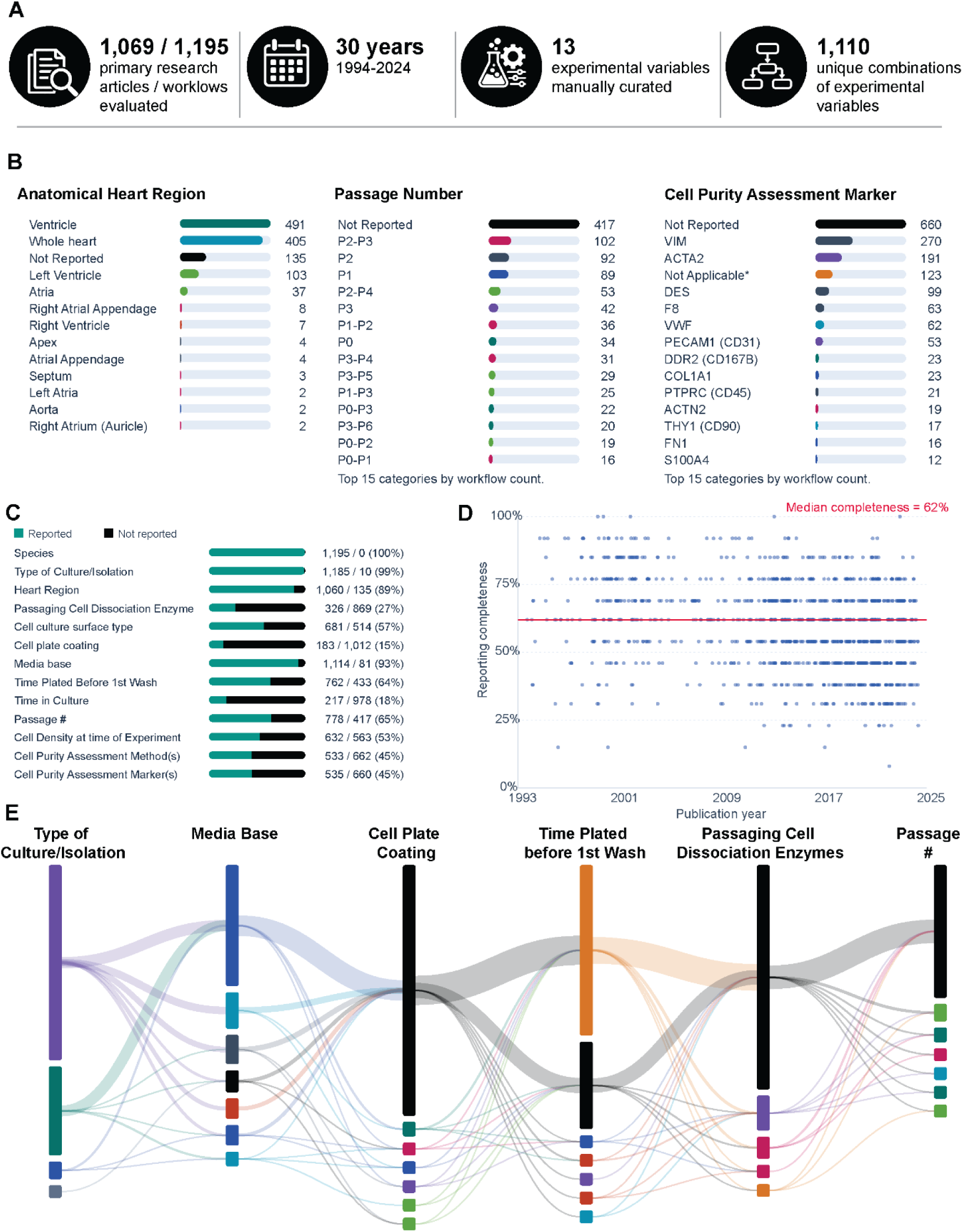
CF culture practices reported across 30 years of literature. A. A review of 1,195 workflows reported among 1,069 primary research publications spanning approximately 30 years was used to evaluate reporting of 13 experimental variables relevant to CF isolation, culture, and characterization. B. Distribution of selected variables illustrating heterogeneity in experimental CF models, anatomical heart region, passage number, and cell purity assessment markers. *Not applicable = When authors used previously isolated or commercially sourced cells and did not describe the assessment method or marker, we coded that field as Not Applicable rather than Not Reported. C. Reporting frequency for experimental variables that define or may influence the cultured CF model. D. Reporting completeness across publication year, calculated for each study as the proportion of applicable experimental variables for which information was reported. E. Sankey diagram of the combinations of six culture variables (cell source, basal medium, plate coating, initial adhesion time, dissociation enzymes, and passage) among studies using human CF.

How cultures were characterized was similarly variable. Fifty-six percent of workflows did not report an assessment of CF identity or culture purity, and those that did used 64 different markers (Figure 7B). Of the 57 markers that are human proteins, we quantified 48, and 33 changed at least 2-fold between P0 and P2 in Substudy I or IV (FDR < 0.05). These included five of the six most frequently used markers: ACTA2, DES, VWF, PECAM1, and F8 decreased, whereas VIM, the most frequently used, changed little. An assessment of CF identity or purity may therefore yield different results across passages. Only 53 studies assessed PECAM1; therefore, PECAM1-positive populations in cultured CF may be under-recognized because they are rarely measured.

Reporting was often incomplete (Figure 7C). Across the 13 variables, 41% of applicable information was not reported. Heart region was not reported in 11% of all studies and 36% of human studies, and 34% of human studies reported ventricle without specifying left or right, although cellular composition and molecular features differ across regions [73]. Reporting completeness changed little over 30 years (Figure 7D), despite increasing emphasis on rigor and reproducibility [11,74–76]. Incomplete reporting also limits the information available to investigators who use previously isolated cells, as in Substudy V.

Among 208 human CF workflows, we identified 140 distinct combinations of cell source, basal medium, plate coating, initial adhesion time, dissociation enzymes, and passage (Figure 7E), with similar diversity in rodent studies (Supplementary Figure 14B). In our substudies, passage alone altered the abundance of hundreds of surface proteins. Diverse methods are not a problem in themselves, but combined with incomplete reporting, they make it difficult to know which culture conditions underlie a given result. The complete literature atlas can be explored in CFxplorer.

### 3.9. CFxplorer integrates an expanded human CF Surfaceome Atlas with molecular and experimental context

CFxplorer brings together three data components from this study (Figure 8): the CF literature atlas, passage-resolved surfaceome and whole cell datasets from Substudies I-V, and a Human CF Surfaceome Atlas. The literature atlas lets users explore how CF have been isolated, cultured, and characterized over time, and the passage-resolved datasets let users follow any protein across passage, donor, tissue handling, and assay.

**Figure 8.**
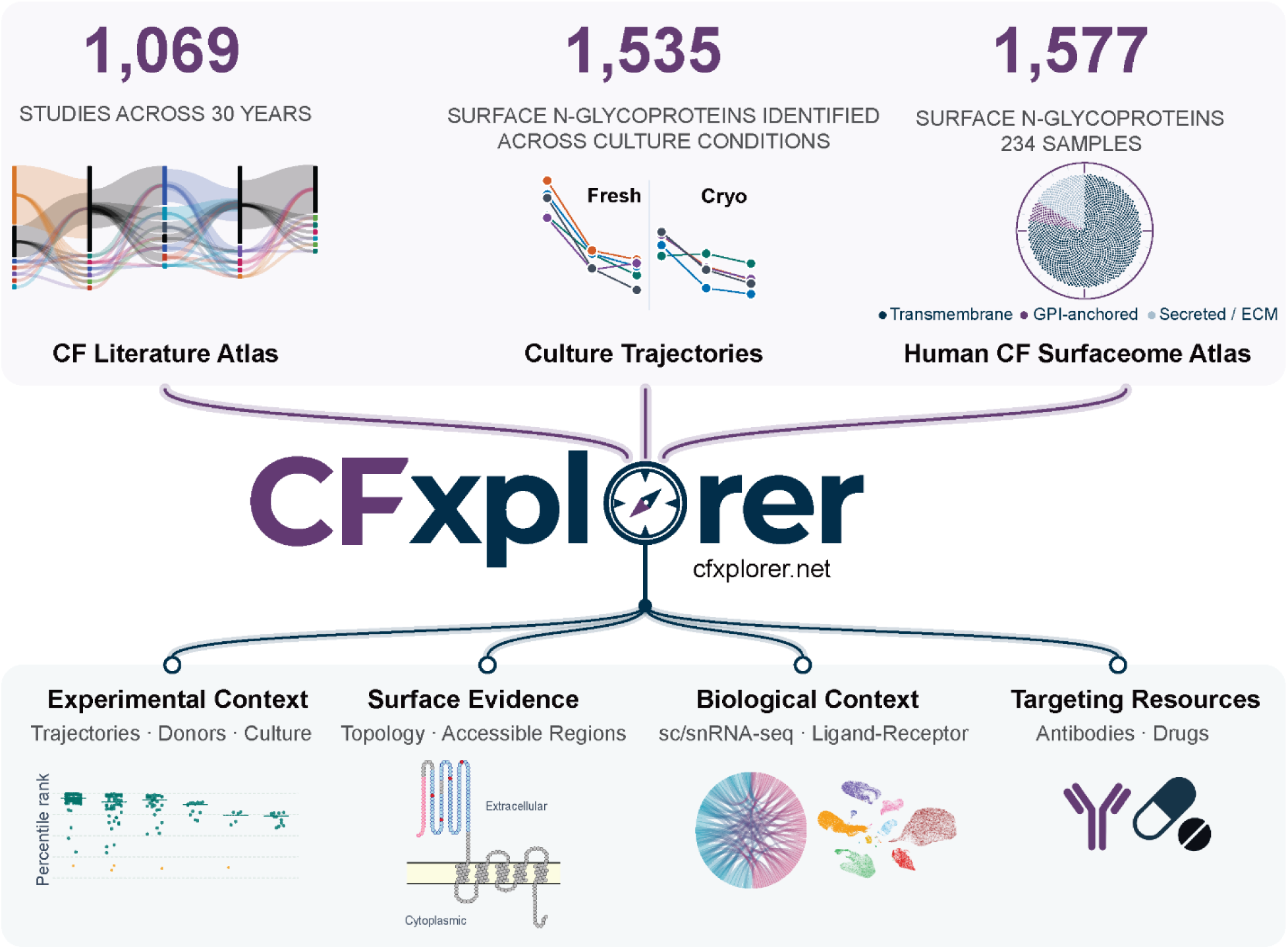
CFxplorer integration of human CF surfaceome data and complementary annotations. CFxplorer brings together CF culture practices and reporting from 1,069 studies spanning 30 years, 1,535 cell surface proteins identified among passage-resolved experimental data from Substudies I-V, and a Human CF Surfaceome Atlas comprising 1,577 cell surface N-glycoproteins identified among 234 samples. The resource integrates these datasets with experimental context, extracellular peptide and topology evidence, cardiac cell-type and ligand-receptor context, and available antibody and pharmacologic resources to support evaluation and prioritization of human CF surface proteins.

To extend the resource beyond the 1,535 surfaceome proteins identified in these experiments, we combined CellSurfer data from 234 human CF samples, including the substudies reported here, CSC method development experiments, and previously published data [26], into an atlas of 1,577 cell surface N-glycoproteins. Experiment-level data preserve detection frequency and peptide spectrum match-based rank across samples, donors, disease states, passages, and methods, so users can view whether, how consistently, and how prominently a protein has been detected in human CF.

Detected peptides and N-glycosylation sites are mapped onto protein topology with Protter [46], providing direct evidence for extracellular regions. Experimental evidence agreed with UniProt-annotated topology for 1,466 proteins and conflicted with it for 37. A further 23 proteins had no resolved topology or localization annotation in UniProt, and 46 were supported by isoform-specific peptides, including three isoforms whose predicted topology differs from that of the canonical protein. Such cases can reveal unrecognized extracellular regions, alternative topologies, or surface-exposed isoforms and support the value of the experimental evidence of surface accessibility provided by CellSurfer [26].

Two human heart sc/snRNA-seq datasets [60,61] are integrated into CFxplorer and show whether transcripts encoding a detected surface protein are enriched in fibroblasts or shared with other cardiac cell types. Curated ligand-receptor pairs, antibody resources, and pharmacologic annotations from ChEMBL and the Broad Institute Drug Repurposing Hub, with links to DrugBank, connect detected proteins with available reagents and approved or investigational compounds. Users can therefore start from a protein of interest or an experimental question and prioritize candidate CF surface markers and targets for validation.

## 4. Discussion

Cultured CF remain essential for studying fibroblast biology and fibrosis, but what they represent depends on how they were generated. Across five sequential substudies, the human CF surfaceome changed most during the first passage, and these changes were reproduced in CF from fresh and cryopreserved tissue. The initial adhesion period altered the abundance of proteins commonly used as endothelial markers; culture shifted CF away from markers of other cardiac cell types and toward an activated, matrix-producing fibroblast state; pharmacologic targets were lost or gained; and commercially sourced CF varied more among donors than with passage. Together with the heterogeneous and incompletely reported culture practices in the literature, these findings support a simple message: the cultures used in a study should be characterized rather than assumed to represent CF in the heart.

### 4.1. Early culture reshapes the CF surfaceome

The largest changes occurred between P0 and P1 in both laboratory-isolated cohorts, and most were shared by CF from fresh and cryopreserved tissue. These findings extend prior studies showing that the fibroblast phenotype is sensitive to culture and passage [18–25] by providing a passage-resolved view of hundreds of extracellularly exposed proteins in primary human CF. In Substudy IV, early culture was associated with increases in proteins of fibroblast activation and matrix production, including FAP, fibrillar collagens, FN1, LOX and ITGA11, consistent with the known effects of serum and stiff culture substrates [18,19]. ACTA2 did not increase, however, and other contractile proteins rose only modestly or not at all, so these cells moved toward a matrix-producing activated state rather than a contractile myofibroblast state over the passages examined. Gene-set analysis supported this: protein secretion, collagen fibril organization and EMT sets increased and oxidative phosphorylation decreased, without a change in canonical TGF-β signaling sets. At the same time, markers that distinguish fibroblasts from other cells in the heart, such as DCN, ABCA6 and CDH19, decreased. Change was not uniform: proteins increased, decreased or remained stable, and the magnitude, and sometimes the direction, of change differed among donors and between preparations. Changes at the cell surface were reflected only in part by whole cell abundance, and each approach measured proteins the other did not, so whole cell proteomics cannot substitute for direct surface measurement when the question concerns surface markers, receptors or antibody-accessible targets. Because P0 already reflects tissue dissociation, adhesion, selection and up to 12 days in culture, differences between native and cultured CF probably extend beyond the trajectories measured here.

### 4.2. The first culture steps set the starting point

Proteins commonly used as endothelial markers were detected in every donor at P0, declined with passage and were more abundant after a 48 h than a 24 h initial adhesion, and a small PECAM1-positive population was detected by flow cytometry. PECAM1 and CD34 also remained detectable in commercially sourced CF at P3-P5. The cellular source of these proteins was not determined. Many proteins are expressed by more than one cell type, antibody-based assessments of cell type specificity have known limitations, and in human heart sc/snRNA-seq data available in CFxplorer, transcripts for several of these proteins, including PECAM1, CD34, CALCRL, and ANGPT2, are detected in subsets of cardiac fibroblasts. These proteins may therefore be present on a subset of CF as well as on other cell types, and we did not use them to infer the cellular composition of the cultures. Whatever their source, the data show that the earliest culture steps, including initial adhesion time, set the starting point for later passages, and that individual markers responded too differently for any single protein to serve as a measure of culture purity.

### 4.3. Differences between failing and non-failing CF at P0

Cultured CF are widely used to compare cells from failing and non-failing hearts. In Substudy II, we found no significant differences in abundance at P0, but 13 proteins, including TNFAIP6, SFRP4, and VCAM1, were detected in at least two of three failing donors and in no non-failing donor, and ECM proteins such as COL1A1, POSTN, COL6A1, and COL6A2 were higher in every failing donor without reaching significance. Differences were therefore present but could not be resolved quantitatively with three donors per group, and they may have diminished during the 12 days before P0, as in canine CF [25]. COL6A1 and COL6A2 were higher in CF from failing hearts at P0 but decreased with passage, thus continued culture could reduce or hide these disease-associated differences. At the same time, culture introduced many changes that could be mistaken for disease effects if failing and non-failing CF are cultured for different lengths of time. Studies comparing cultured CF from failing and non-failing hearts should report culture history and confirm key differences in minimally cultured or in situ CF.

### 4.4. CF markers and pharmacologic targets change with culture

Proteins commonly used to identify or characterize CF increased with passage in Substudy IV, including FAP, THY1, and COL1A1 in both arms and CD44, FN1, and PDGFRA in the fresh arm, while pharmacologic targets were lost (AOC3, ANGPT2, ASAH1) or gained (EGFR, NOTCH2, NPR3, HAPLN1). AOC3 illustrates the point: its surface abundance fell by a median of more than 10-fold between P0 and P2 in Substudies I and IV and was higher after a longer initial adhesion. Rising marker levels therefore did not show that cultured CF kept the profile of fibroblasts in the heart, because markers that define fibroblasts in the human heart atlas decreased over the same passages (Section 3.7). Conventional markers remain useful, but a culture can retain CF-associated proteins while much of its surfaceome changes. VIM, the marker most often used in the literature to assess CF identity, is an example. Its whole cell abundance changed by less than 1.3-fold between P0 and P2, whereas five of the six most frequently used markers and hundreds of other proteins changed substantially over the same passages. Detection of a stable marker therefore shows that a culture retains a feature expected of CF, not that the cells are unchanged. VIM is also expressed by several other non-myocyte cell types in the heart kane, so its detection alone does not confirm that a culture consists of CF. Investigators should therefore characterize their cultures at the passage used for experiments and measure directly any surface protein central to their study.

### 4.5. Commercially sourced cells and donor variation

Commercially sourced CF gave a different picture than cells isolated in our laboratory. Although all five donors were from non-failing hearts, their cells differed considerably from one another from the first passage studied: donor accounted for about half of the variance in a typical protein, and the surface profiles of different donors were less alike at P3 than those of Substudy IV donors at P0. Across P3-P5, protein abundance changed within individual donors, but the changes differed in direction and magnitude among donors, so few reached significance at the cohort level. Some proteins nevertheless changed in the same direction in most donors: ISLR, PLAU, and CADM1 changed at the cell surface in every donor, the pharmacologic targets NOTCH2 and HAPLN1 increased at the surface in four of five donors, as they did in laboratory-isolated cells, and calponin (CNN1) decreased in every donor in whole cell data. Across the whole cell proteome, 51% of proteins changed in the same direction in at least four of five donors, more than the 37.5% expected by chance, whereas at the cell surface the proportion (37%) did not exceed chance. Because these cells were purchased and we could not obtain their isolation or culture history, we cannot tell whether this variation reflects the donors themselves or how the cells were handled before P3. Nor can we separate commercial source from the later passages at which these cells were studied. Whatever the cause, averaging across donors can obscure changes that are real within individuals. Studies using purchased cells should treat donors, or lot, as a major source of variation, report lot and passage history where available, and examine changes within donors as well as across them. For this reason, our analyses emphasize within-donor change, and CFxplorer preserves donor-level data.

### 4.6. Reporting culture history

Several of the variables that affected the surfaceome in our substudies differed widely across published studies or were not reported. Passage ranged from P0 to P39, many studies used mixed or unreported passages, and 41% of information across all 13 variables was not reported, with little improvement over 30 years. Diverse methods are not a problem when conditions are clearly reported, and different questions may require different approaches. Incomplete reporting, however, makes it difficult to know whether differences among studies reflect biology, culture history, or both. We therefore suggest that studies report, at minimum, cell source, passage, initial adhesion time, tissue handling, and how and at which passage culture composition was assessed.

### 4.7. CFxplorer as a resource for investigator-defined questions

Because CF cultures differ in how they are generated, and individual proteins differ in how they respond to culture, investigators need a way to check how proteins relevant to their own work behave across these conditions. CFxplorer was developed for this purpose. Rather than reducing the data to a few representative proteins, the dashboard allows investigators to start from a protein or an experimental question and view whether a protein is stable or passage-sensitive, whether its trajectory is shared across donors and preparations, and whether its surface representation parallels whole cell abundance. The Human CF Surfaceome Atlas provides direct evidence of extracellular accessibility that annotation-based predictions can miss, including extracellular regions that conflict with annotated topology and surface-exposed isoforms. These data do not establish any protein as a validated CF marker or therapeutic target, but they can help investigators design experiments and choose candidates for validation.

### 4.8. Limitations

Each substudy included few donors, so the data describe culture-associated changes rather than population-level reference ranges, and Substudy II could detect only large differences in abundance. Because the substudies used two CellSurfer workflows and two MS instruments, comparisons across substudies concerned the direction and pattern of change rather than abundance. Proteins near the detection limit were more often missing and were not imputed; thus changes in low-abundance proteins may be under-represented. Bulk measurements also cannot distinguish changes within CF from selection among CF subpopulations or changes in other cell types. Finally, our analyses describe protein abundance rather than function: the signature, gene-set, and ligand–receptor analyses do not measure pathway activity or signaling, the atlas signatures compare cultured cells with transcript markers rather than with the proteome of CF in the heart, and fibroblast activation was not assessed by functional measures such as contraction or collagen deposition.

Our study also examined only some of the variables that differ among CF culture systems. In Substudy V, commercial source and later passage cannot be separated, and in all substudies, cells were grown on two-dimensional culture surfaces. Hydrogel scaffolds that reproduce the mechanical and biochemical properties of the cardiac ECM now allow fibroblasts to be cultured in 2D, 2.5D, and 3D formats [77–80], and 2.5D models have been applied successfully to a range of cell types [80–82]. Whether these systems preserve the surfaceome of CF at P0 more closely than conventional culture is an important question for future work. Our data therefore do not define an optimal culture method, but they provide a baseline against which such models can be compared.

### 4.9. Conclusion

Culture history is a defining feature of the human CF model. The CF surfaceome changes most in the first passage, the earliest culture steps influence the proteins present at P0, markers, targets, and matrix programs move with culture, and commercially sourced cells vary considerably among donors. Together with the heterogeneous and incompletely reported practices in the literature, these findings argue for characterizing cultured CF rather than assuming what they represent. CFxplorer makes the underlying data available so that investigators can evaluate the proteins relevant to their own questions and prioritize candidate markers and targets for validation. We urge researchers to consider culture history when designing and interpreting studies of cultured CF, and to report culture conditions in sufficient detail that results can be compared and reproduced.

## Data Availability

MS data files are available at MassIVE (https://massive.ucsd.edu/ProteoSAFe/; accession: MSV000103688; password: cf2026).

## Acknowledgements

This work was supported by the National Institutes of Health [R35-HL155460] and the Omaha Community Foundation to RLG, American Heart Association [20PRE35200049] to LBL, and National Institutes of Health [1F31HL189247] to ER. Funding sources were not involved in study design, data collection, interpretation, analysis, or publication. We are thankful to Walter Bussey, Blake Hoffmeyer, Kristine Spears, Ryan Ruskamp, Dr. Stan Radio, Dr. Anthony Castleberry, and Dr. John Um for assistance with specimen collections, and Shelby Konfrst, Michelle Waknitz, and Chase Castro for assistance with cell culture and imaging. Specimens and data used in this study were obtained from the Medical College of Wisconsin and the Nebraska Cardiovascular Biobank and Registry, which is supported by the UNMC Center for Heart and Vascular Research and the Omaha Community Foundation. Analyses were performed using instrumentation in the UNMC Flow Cytometry Research Facility (RRID:SCR_017736), administrated through the Office of the Vice Chancellor for Research, and supported by state funds from the Nebraska Research Initiative and The Fred and Pamela Buffett Cancer Center’s National Cancer Institute Cancer Support Grant. Major instrumentation has been provided by the Office of the Vice Chancellor for Research, The University of Nebraska Foundation, the Nebraska Banker’s Fund, and by the NIH-NCRR Shared Instrument Program. We acknowledge use of the UNMC Advanced Microscopy Core Facility, RRID:SCR_022467, supported by the National Institutes of Health, Fred and Pamela Buffett Cancer, Ne-INBRE, Nebraska Research Initiative, and UNMC Vice Chancellor for Research Office; CardiOmics Program supported by the UNMC Center for Heart and Vascular Research; UNMC Multiomics Mass Spectrometry Core Facility (RRID: SCR_012539); and the Bioassay Core (RRID:SCR_026270) supported by the NIH-funded Center for Heart and Vascular Research Centers of Biomedical Research Excellence (COBRE).

## Author Contributions

**Roneldine Mesidor:** Data curation, Formal analysis, Investigation, Methodology, Validation, Visualization, Writing – review and editing. **Linda Berg Luecke:** Conceptualization, Data curation, Formal analysis, Funding acquisition, Investigation, Methodology, Validation, Visualization, Writing - original draft, Writing - review and editing. **Matthew Waas:** Data curation, Formal analysis, Investigation, Methodology, Software, Validation, Visualization, Writing - review and editing. **Jennifer Schneider:** Data curation, Formal analysis, Validation, Visualization, Writing - review and editing. **Garam Park:** Data curation, Investigation, Methodology, Writing - review and editing. **Ella Rowe:** Data curation, Formal analysis, Funding acquisition, Investigation, Writing – review and editing. **Kyu Shik Kim:** Data curation, Formal analysis, Investigation, Visualization, Writing - review and editing. **Erin N. Schuette:** Investigation, Methodology, Writing - review and editing. **Pranav Renavikar:** Formal analysis, Investigation, Validation, Writing - review and editing. **Heather C. Jensen-Smith:** Formal analysis, Investigation, Validation, Visualization, Writing - review and editing. **Claudius Mahr:** Investigation, Methodology, Writing - review and editing. **Daniel R. Anderson:** Investigation, Methodology, Resources, Writing - review and editing. **Rebekah L. Gundry:** Conceptualization, Data curation, Formal analysis, Funding acquisition, Investigation, Project administration, Resources, Software, Supervision, Validation, Visualization, Writing - original draft, Writing – review and editing.

## Declaration of generative AI use

Generative AI tools were used to assist with programming and development of CFxplorer.net and for grammatical editing of the manuscript. AI was not used to generate experimental data, perform scientific calculations or statistical analyses, or independently interpret results. All scientific analyses, interpretations, and conclusions were performed or verified by the authors.

## Disclosures

R.L.G. is on the advisory board of ProtiFi, LLC and receives no compensation of any kind for this role. The remaining authors declare no competing interests.

## Notes

https://cfxplorer.net

https://massive.ucsd.edu/ProteoSAFe/

