## Supplementary Methods and Figures for "Culture-Associated Changes in the Human Cardiac Fibroblast Surfaceome: A Proteomic Resource for Cell Surface Marker and Target Discovery"

### These authors contributed equally

**Supplementary Methods**

*Imaging*

Fibroblast morphology was assessed by brightfield and widefield microscopy. In Substudy I, immunocytochemistry for vimentin (fibroblast marker) was used, whereas Substudies IV-V assessed vimentin, α–smooth muscle actin (α-SMA; activated fibroblasts/myofibroblasts), wheat germ agglutinin (WGA; cell membrane), and DAPI (DNA). For Substudy I, cells were plated on coverslips and images were obtained with a Zeiss 710 Laser Scanning Microscope. For Substudies IV-V, cells were plated on 4-well chamber slides and whole slide images were obtained with the Zeiss Axioscan7 whole slide imaging system using a 40x (0.95 NA) objective and identical data collection parameters.

For Substudies I, II, and IV, histological analysis of hematoxylin and eosin and Masson's trichrome of tissue sections was performed. At the time of collection, tissue pieces were fixed in 10% neutral buffered formalin at room temperature for 24 to 48 hours. After fixation, formalin was removed and replaced with 70% ethanol for storage. Once all tissues were collected, they were provided to the Tissue Sciences Facility at UNMC for paraffin embedding, sectioning, and staining. Briefly, sections were deparaffinized in xylene and rehydrated using graded alcohols. We performed hematoxylin and eosin (H&E) staining on an automated stainer and Masson's trichrome staining using the Masson's Trichrome Stain Kit (StatLab, KTMTR) according to the manufacturer's protocol. Images were collected and evaluated by a pathologist to assess the degree of fibrosis in the tissue used for CF isolation. All methodological details are described in Supplementary Table 2.

*Immunofluorescence of explanted CF*

Isolated CF were cultured on coverslips were washed twice with $DPBS^{-/-}$. Cells were fixed for 5 minutes in a fixation solution (4% PFA in PBS) at 4 °C. Cells were washed twice with 2 mL of PBS in a 6-well plate. Cells were permeabilized for 5 minutes in a permeabilization solution (0.1% Triton-X in 0.1% BSA in PBS) and subsequently blocked for 60 minutes in a blocking solution (1% BSA, 10% goat serum, 0.3 M glycine, 0.1% Tween in PBS). The primary antibody was diluted in blocking solution and added to coverslips in a humidified chamber for overnight incubation at 4 °C. Antibody details are provided in Supplementary Table 2. Next, cells were washed twice with PBS. Cells were incubated with Hoechst 33342 for 10 min at room temperature in the dark. After incubation, the cells were washed twice with PBS. Finally, 10 µL of mounting medium (Prolong Diamond Antifade Mountant) was added to the coverslips. Images were captured using a Zeiss 710 Laser Scanning Microscope. ZEN imaging software was used to process images.

Isolated CF cultured on chamber slides were washed three times with cold $DPBS^{+/+}$. Cells were fixed for 15 minutes in fixation solution (4% PFA in $DPBS^{+/+}$) at room temperature. Cells were washed three times with cold $DPBS^{+/+}$ and stored in a humidified chamber at 4 °C. Once all slides were obtained for each donor, slides were collected and incubated with WGA for 10 minutes at room temperature. Next, cells were washed three times with cold $DPBS^{+/+}$ and subsequently permeabilized for 5 minutes in permeabilization solution (0.1% Triton X-100 in $DPBS^{+/+}$). After incubation, cells were washed thrice with cold $DPBS^{+/+}$ and subsequently blocked for 60 minutes in blocking solution (1% BSA, 10% goat serum, 0.3 M glycine, 0.1% Tween in $DPBS^{+/+}$). Primary antibody was diluted in blocking solution and added to slides placed in a humidified chamber for an overnight incubation at 4 °C. Antibody details are provided in Supplementary Table 2. Next, cells were washed three times with cold $DPBS^{+/+}$. Secondary antibody was diluted in blocking solution and added to appropriate slides for 60 minutes. After the incubation, cells were washed three times with cold $DPBS^{+/+}$. Finally, mounting medium was added to each slide, and coverslips were added to each slide. Slides were cured for 24 h at room temperature in the dark and imaged with a Zeiss Axioscan7 whole slide imaging system. QuPath version 0.5.1 [1] was used to segment and quantitatively assess fluorescence using identical analysis parameters.

*Microscale Cell Surface Capture (μCSC)*

All steps were performed on ice, unless otherwise noted, and all solutions were prepared fresh on the day of the labelling, as previously described [2]. Cells were labeled while adherent to the plate. Cells were washed twice with room temperature wash solution (1X PBS, 0.1%FBS) and once with cold labeling buffer (1X PBS, 0.1% FBS, pH 6.0) before oxidization with 1 mM sodium-meta-periodate in cold labeling buffer for 15 min in the dark at 4 °C. Oxidation solution was removed, quench solution (2 mM sodium sulfite in cold labeling buffer) was added to cells and incubated for 10 minutes in the dark at 4 °C. Cells were washed twice with cold labelling buffer, followed by incubation in 10 mM biocytin hydrazide and 50 mM (1H-imidazol-2-Yl)-methanamine dihydrochloride in cold labeling buffer for 60 min in the dark at 4 °C. Cells were washed three times with cold wash buffer (1X PBS, 0.1% FBS). Cells were resuspended in lysis buffer (40% 250 mM ammonium bicarbonate/ 40% 5X invitrosol/20% 100% acetonitrile) and scraped off the plate. Cells were sonicated using a VialTweeter with three intervals of ten-second pulses with ten seconds of cooling on ice between repetitions. Proteins were reduced with 5 mM tris(2-carboxyethyl)phosphine (TCEP) for 30 min at 37 °C with vortexing, followed by alkylation with 10 mM 2-iodoacetamide (IAA) for 30 min at 37 °C with vortexing. Protein quantitation was determined using the Pierce 660 nm Protein Assay Reagent. Trypsin/Lys-C mixture was added at a 1:40 enzyme-to-substrate ratio; pH was adjusted to ~8.5 by addition of 2N NaOH and digested overnight at 37 °C with vortexing. All vortexing was performed in a thermomixer at 1200 rpm. Peptide quantitation was performed using Pierce Quantitative Fluorometric Peptide Assay. 135-235 μg of peptide was diluted in binding buffer (80 mM sodium phosphate, 2 M NaCl, 0.2% Tween 20, 25 μM EDTA, 0.1% SDS, pH 7.8) and boiled for 5 minutes at 95 °C in a heat block, followed by cooling on ice for 5 minutes. 100 μL of GenScript Streptavidin MagBeads (GenScript, L00424) were equilibrated in binding buffer (80 mM sodium phosphate, 2 M NaCl, 0.2% Tween 20, pH 7.8) and added to the peptides and incubated for 1 h with mixing. Beads were washed sequentially with: (1) 2% SDS in 1× PBS, (2) 80 mM sodium phosphate, 2 M NaCl, 0.2% Tween 20, pH 7.8, (3) 100 mM sodium carbonate, and (4) 50 mM ammonium bicarbonate. Peptides were released by incubating the beads in 40 μL 50 mM ammonium bicarbonate containing 2 μL PNGase F overnight at 37 °C with vortexing. Supernatant was removed to a new tube, beads were rinsed once with 20 μL 0.1% SDS in ultrapure water, and the rinse was combined with the supernatant. Peptides were cleaned using SP2 to remove detergents and other contaminants, as previously described [3]. Glycopeptide capture and bead washing were performed using an epMotion 5073m (Eppendorf).

*Nanoscale Cell Surface Capture (nanoCSC)*

All steps were performed on ice, unless otherwise noted, and all solutions were freshly prepared on the day of labelling. Cells were labeled while adherent to the plate. Cells were washed twice with room-temperature wash solution (1X PBS, 1% FBS) and once with cold labeling buffer (1X PBS, 1% FBS, pH 6.5) before oxidization with 1.6 mM sodium metaperiodate in cold labeling buffer for 15 min in the dark at 4 °C. Oxidation solution was removed, and cells were washed twice with cold labelling buffer, followed by incubation in 0.5 mM Alkoxyamine-PEG4-Biotin and 5 mM 5-methoxy-anthranilic acid in cold labeling buffer for 30 min in the dark at 4 °C. Cells were washed three times with cold labeling buffer. Cells were lysed in 9.5% sodium dodecyl sulfate (SDS), 50 mM triethylammonium bicarbonate (TEAB), 0.01% n-dodecyl β-D-maltoside (DDM), 2mM magnesium chloride ($MgCl_2$), pH 7.55 with 85% phosphoric acid and 250 units of Benzonase, and scraped off the plate. Cells were sonicated using a VialTweeter with three intervals of ten-second pulses with ten seconds cooling on ice between repetitions. Protein quantitation was determined using the Pierce 660 nm Protein Assay Reagent and a Varioskan LUX Multimode Microplate Reader with SkanIt 5.0 software (Thermo Fisher Scientific) per manufacturer instructions. Proteins were processed using a suspensiontrapping (S-trap™) protocol on an OT2 robot (Opentrons) as described [4]. Briefly, proteins were reduced with 5 mM tris(2-carboxyethyl)phosphine (TCEP) for 10 min at 55 °C with vortexing, followed by alkylation with 10 mM 2-iodoacetamide (IAA) for 10 min at 37 °C with vortexing. Samples were acidified with 27.5% phosphoric acid and then diluted with S-Trap™ binding buffer (100 mM TEAB, 90% methanol with pH 7.1). Samples were loaded onto an S-Trap™ 96-well mini plate and washed three times with S-Trap™ binding buffer and three times with 2:1 chloroform: methanol. Trypsin-LysC in 50 mM TEAB solution was added to the S-Trap™ at a ratio of 1:15 protease: protein. The S-Trap™ was incubated at 37 °C for 2 h. Peptides were eluted sequentially with (1) 50 mM TEAB, 0.02% DDM, (2) 0.2% aqueous formic acid, 0.02% DDM, and (3) 50% acetonitrile (MeCN), 0.2% formic acid, 0.02% DDM. S-trap™ processing was performed using an Opentrons OT2 liquid handler.

Peptide samples were measured using the Pierce Quantitative Fluorometric Peptide Assay and a Varioskan LUX multimode microplate reader with SkanIt 5.0 software (Thermo Fisher Scientific), following the manufacturer's instructions. Next, 35-62 μg of peptide was diluted in binding buffer (80 mM sodium phosphate, 2 M Sodium chloride (NaCl), 0.2% Tween 20, 25 μM EDTA, 0.1% SDS, 0.2% DDM, pH 7.56) and 1.25X Halt™ Protease and Phosphatase Inhibitor Cocktail to inhibit residual trypsin activity. Next, 10 μL of GenScript Streptavidin MagBeads (GenScript, L00936) were equilibrated in binding buffer (80 mM sodium phosphate, 2 M NaCl, 0.2% Tween 20, 0.2% DDM, pH 7.8) and added to the peptides and incubated for 30 min with mixing. Beads were washed sequentially with (1) 2% SDS in 1X PBS, 0.2% DDM, (2) 80 mM sodium phosphate, 2 M NaCl, 0.2% Tween 20, 0.2% DDM, pH 7.56, (3) 100 mM sodium carbonate, 0.2% DDM, and (4) 50 mM ammonium bicarbonate, 0.2% DDM. Peptides were released by incubating the beads in 30 μL of 50 mM ammonium bicarbonate containing 2 μg PNGase F at 37 °C with vortexing for 1h. Supernatant was removed to a new tube; beads were rinsed once with 10 μL 0.1% SDS, 0.1% DDM in LC-MS water, and the rinse was combined with the supernatant. Peptides were cleaned using SP2 to remove detergents and other contaminants as previously described [3]. Glycopeptide capture, bead washing, and peptide cleanup were performed using an Opentrons OT2 liquid handler.

*Whole cell proteomics*

Unless otherwise noted, all chemicals were obtained from Sigma-Aldrich. S-trap™ Turbo plate (Protifi) was used for sample processing on an OT2 robot (OpenTrons). Briefly, samples were lysed in 10% sodium dodecyl sulfate, 50mM triethylammonium bicarbonate (TEAB), 2 mM magnesium chloride, and adjusted to pH 7.55 with 85% phosphoric acid. 250 units of benzonase was added and incubated at room temperature for 10 minutes. Following protein quantitation by the 660 nm method kit (Pierce), 100 µg of each sample was reduced with 10 mM TCEP in a heater-shaker module for 15 minutes set at 55°C at 1200 rpm, followed by alkylation with 20 mM iodoacetamide (IAA) for 30 minutes in the dark at room temperature at 1200 rpm. Next, 27.5% phosphoric acid was used to acidify the proteins, and the samples were diluted with 300 µL of S-Trap™ binding buffer (100 mM TEAB, 90% methanol, pH 7.1). The samples were then loaded onto an S-Trap™ Turbo plate and washed three times with 2:1 chloroform: methanol and then three times with S-Trap™ binding buffer. 8 µg of Trypsin-LysC in 50 mM TEAB solution was added to the top of the S-Trap™ Turbo and then incubated at 37°C for 2 hours. Peptides were eluted serially with 50 mM TEAB and 0.4% aqueous formic acid with 0.04% n-Dodecyl β-D-maltoside (DDM, GoldBio), then dried by vacuum centrifugation. Peptide samples were diluted to 0.2 µg/µL and measured using a Pierce Quantitative Fluorometric Peptide Assay (Fisher Scientific) using a Varioskan LUX multimode microplate reader (Thermo Fisher).

*MS data acquisition*

In substudies I-III, samples were analyzed using a Dionex UltiMate™ 3000 RSLCnano liquid chromatography (LC) system. They were separated using a PepMap RSLC C18 (substudies I-II) and a µPac™ Neo 50 cm (substudy III), both coupled to an Orbitrap Exploris™ 480 mass spectrometer (Thermo Fisher Scientific). In substudies IV-V, samples were injected onto a Vanquish™ NEO liquid chromatography (UPLC) system and separated using an IonOpticks Aurora® Ultimate™ C18 for nanoCell surface capture and a µPAC™ Neo 50 cm column for Whole Cell capture, coupled to an Orbitrap Astral mass spectrometer (Thermo Fisher Scientific). Gradient, data acquisition, and analysis settings are listed in the supplementary information (Supplementary Table 4). Sample sets spiked with an iRT synthetic peptide mixture were blocked and acquired in a randomized order, with digested E. coli injected immediately before and after every sixth sample to evaluate system performance. The consistency and reproducibility of the system performance were evaluated using QuiC software (5.3.240326.57353).

*MS data search*

Raw files were analyzed using Spectronaut™ 21 (Biognosys, Switzerland) using directDIA+™. Cell surface capture (CSC) and whole cell lysate datasets were analyzed separately. Spectra were searched against the human Uniprot/SwissProt reference database, including isoforms (42,438 sequence entries, June 2023) and the porcine trypsin reference database (1, December 2019). CSC searches additionally included the PNGase F reference database (1, December 2019) and the streptavidin reference database (1, December 2019). The search and validation parameters, including enzyme specificity, modifications, and mass tolerance, are listed in Supplementary Table 4.

*MS Data processing and protein quantification*

Precursor-level reports (one row per precursor per run) were exported from Spectronaut without cross-run normalization and without imputation, so that they contained the measured values only. Protein quantities were then calculated in R rather than taken from the software's protein report, so that the same procedure applied to every dataset and no protein group was removed by software-specific quantification rules. Precursors were retained in each run if they passed a precursor Q-value of 0.01 and a protein Q-value of 0.01, with a posterior error probability of at most 0.01, matching the cutoffs used in the search. Each precursor was kept in the protein group assigned by the software's protein inference (unique and razor peptides (search software protein inference)). Precursor intensities were summarized to one value per protein group and run with the MaxLFQ algorithm [5], as implemented in the iq R package [6] (version 2.0.1), without normalization or imputation; normalization and missing values were handled in the subsequent steps described below. A protein group was considered identified if at least one of its precursors passed these cutoffs in at least one sample; a protein group was considered quantified, and was tested, if it was observed in enough samples to meet the eligibility rule given below.

CSC protein groups were classified with Veneer [7] after protein quantification, and only those classified as High or Medium confidence cell surface proteins were retained. Repeat injections of the same biological sample were combined by taking the median per protein, before any statistical analysis. The following samples were excluded before analysis: Subtudy IV CSC Fresh donor 1376 P0 (Known failed acquisition).

*MS data normalization and missing values*

Normalization and missing-value handling were chosen empirically for each dataset before the final analysis, by passing every candidate combination through the full statistical model [8]. Each combination was evaluated for false positives in data with the condition labels shuffled within donor (40 shuffles per combination), and for sensitivity and false discovery proportion in simulated data built from the real data with known changes added (10 simulations per scenario). Whether each dataset's sample-wide abundance shift followed the biological condition was also tested, because normalization removes such a shift. Imputation was considered only if it increased sensitivity substantially without increasing false positives.

The final analysis used no normalization for Substudy III CSC; median normalization (each sample's median, over proteins quantified in every sample, aligned to the common median) for Substudy I CSC, Substudy II CSC, Substudy IV CSC, Substudy IV WCL, Substudy V CSC and Substudy V WCL. No missing values were imputed; proteins were analysed using the observed values only. The locked choice differed from the per-dataset diagnostic recommendation for Substudy I CSC, Substudy IV CSC (Fresh) and Substudy IV WCL (Cryo), to apply one method consistently across datasets addressing the same question. Substudy III CSC was analysed without normalization because the diagnostic showed that the sample-wide shift followed the biological condition consistently across donors, so normalization would have removed a biological difference. For each dataset, the sample-wide shift was quantified before normalization as the change in each sample's median abundance over proteins quantified in every sample and tested against the condition with donor as a blocking factor.

Missing values were assessed at the protein level for each dataset, on the values as analyzed and before any imputation. Between 1.6 and 27.3% of values were missing across datasets. The proportion of missing values per protein was compared with its mean observed abundance (Spearman rho from -0.75 to -0.25); missingness was concentrated at low abundance in Substudy I CSC, Substudy II CSC, Substudy III CSC, Substudy IV CSC (Fresh), Substudy IV CSC (Cryo), Substudy IV WCL (Cryo), Substudy V CSC and Substudy V WCL. Proteins were filtered by the eligibility rule before any imputation would be applied, and no values were imputed, so proteins were tested on observed values only. Missingness associated with the condition was screened per protein with Fisher's exact test (Benjamini-Hochberg correction); proteins detected in only one condition were reported as detected or not detected rather than assigned imputed fold changes.

*Statistical analysis*

Differential abundance was assessed per dataset with linear models in limma [8,9], with donor as a blocking factor where donors were measured in more than one condition. Conditions and the requirement for a protein to be tested, per dataset: Substudy I CSC (P0, P1, P2; tested if at least 2 donors observed at every passage); Substudy II CSC (non-failing,failing; tested if at least 3 donors observed in each group); Substudy III CSC (T24, T48; tested if at least 4 donors observed at each condition); Substudy IV CSC (Fresh) (P0, P1, P2; tested if at least 3 donors observed at every passage); Substudy IV CSC (Cryo) (P0, P1, P2; tested if at least 3 donors observed at every passage); Substudy IV WCL (Fresh) (P0, P1, P2; tested if at least 3 donors observed at every passage); Substudy IV WCL (Cryo) (P0, P1, P2; tested if at least 3 donors observed at every passage); Substudy V CSC (P3, P4, P5; tested if at least 3 donors observed at every passage); Substudy V WCL (P3, P4, P5; tested if at least 3 donors observed at every passage). Empirical Bayes moderation used an intensity-dependent prior variance trend and robust hyperparameter estimation [9–11] (limma eBayes(trend=TRUE, robust=TRUE)). The primary test for each protein was the moderated F-test across all conditions, with Benjamini-Hochberg correction [10] across all tested proteins in that dataset. Proteins with FDR < 0.05 were reported as primary evidence and those with 0.05 <= FDR < 0.10 as secondary evidence. Pairwise log2 fold changes, nominal P values and pairwise FDR are reported as supporting results.

As a sensitivity analysis, the same models were refitted with DEqMS [12], which estimates each protein's prior variance from the number of peptides used to quantify it. Peptide counts were taken from the same precursor reports.

*Concordance between tissue preparations*

To test whether passage-associated changes defined in one arm (cryopreserved tissue) were reproduced in the other (fresh tissue) from the same donors, the proteins significant in the discovery arm (FDR < 0.05) were evaluated in the replication arm by (i) the proportion changing in the same direction, (ii) the Spearman correlation of fold changes across all shared proteins and (iii) a passage score built from the discovery-arm proteins and applied to each replication-arm sample. Significance was assessed by permutation (1,000 permutations), using replication-arm data from which the passage effect had been removed, so that the null distribution preserved the correlation structure between proteins. Each test was repeated after removing each sample's median, to confirm that agreement did not depend on the sample-wide shift. Differences between arms were tested with a joint model including an arm-by-condition interaction, reported as supplementary because of limited power.

*Visualization and descriptive analyses*

Principal component analysis used the log2 abundances as analyzed (after the locked normalization), restricted to proteins quantified in every sample of the plot, centered but not scaled. Volcano plots show the pairwise $\log_2$ fold change against the nominal P value; proteins detected in one condition only (in at least one donor of that condition and no donor of the other) are shown separately, by the number of donors in which they were detected, because no fold change can be estimated for them. Coordination of the passage response among donors was summarized as the Spearman correlation, for each pair of donors, of their within-donor changes between the first and last condition. The timing of change was assessed by comparing each protein's median within-donor change over the early interval with that over the late interval. Agreement between datasets and between assays was summarized by the Spearman correlation and the proportion of shared proteins changing in the same direction, using median within-donor changes between the first and last condition. Datasets without shared donors across conditions (failing vs non-failing hearts) were compared with the passage datasets through their limma log2 fold changes, for proteins tested in both (matched on leading UniProt accession): Spearman correlation across all shared proteins, and the proportion of proteins significant in the passage dataset (FDR < 0.05) whose fold change had the same sign in the failing vs non-failing comparison. For selected markers and pharmacologic targets, within-donor changes between the first and last condition were plotted per donor; proteins detected at only one endpoint are shown as gain or loss of detection rather than given an imputed value. These analyses are descriptive and were not used to declare significance.

*Trajectory clustering*

Proteins were grouped by the shape of their abundance trajectory with fuzzy c-means clustering, following the Mfuzz approach [13,14] and implemented with the e1071 R package. For each protein, the trajectory was the median, across donors observed at every condition, of the within-donor change from the first condition (no imputation), standardized to mean 0 and standard deviation 1. All proteins meeting this requirement were clustered, regardless of statistical significance; significance from the differential abundance analysis is shown within each cluster. The fuzzifier m was estimated from the number of proteins and conditions [15]. To choose the number of clusters, the minimum distance between cluster centroids was computed for k = 2-15 (20 runs each). Because standardized three-condition trajectories lie on a circle, the distribution of trajectory shapes was also compared with trajectories recomputed after shuffling the condition labels within each protein and donor, and shapes occurring more often than in the shuffled data were taken to indicate distinct trends. Cluster reproducibility was assessed as the mean Jaccard similarity of each cluster across 50 bootstrap resamples of proteins. Clusters were: Substudy I CSC, k = 3. Clustering is descriptive.

*Live cell flow cytometry for CD31 immunodetection*

To generate single-cell suspensions of CF for flow cytometry in substudy I, cells were washed once with Dulbecco's phosphate-buffered saline without calcium and magnesium ($DPBS^{-/-}$) and were treated with TrypLE for 5 minutes at 37 °C. Following collection, cells were washed once with DPBS-/-, counted (trypan blue staining, hemocytometer), and 500,000 cells per sample were transferred into a 96-well plate. CD31 was assessed by flow cytometry as described in Supplementary Table 2.

CD31 immunodetection was assessed at days 15 (passage 0), 20 (passage 1), and 25 (passage 2) of donor 3 (Supplementary Table 2). All steps were performed on ice or at 4 °C. Cells were blocked with 2% FBS in $HBSS^{-/-}$ (block solution) for 15 min. Primary antibodies were added to the cells and gently agitated on an orbital shaker for 45 min. Antibody details are provided in Supplementary Table 2. Cells were washed three times with wash solution (0.5% FBS in $HBSS^{-/-}$). Sytox blue viability stain was pre-diluted in wash solution, used to resuspend cells post-wash, and then gently agitated on an orbital shaker for 5 min. Cells were resuspended in wash solution and transferred to filter-cap flow tubes for analysis by flow cytometer. For all live-cell flow, 10,000 events were collected. Gating of cells was performed with SSC-H x FSC-H on an Attune Nxt flow cytometer (Thermo Fisher Scientific). Single cells were gated using sequential gates of FSC-W x FSC-H and then SSC-W x SSC-H. Live cells were gated using SSC-H x Sytox blue. FlowJo (version 10.7.2) was used to analyze flow cytometry data.

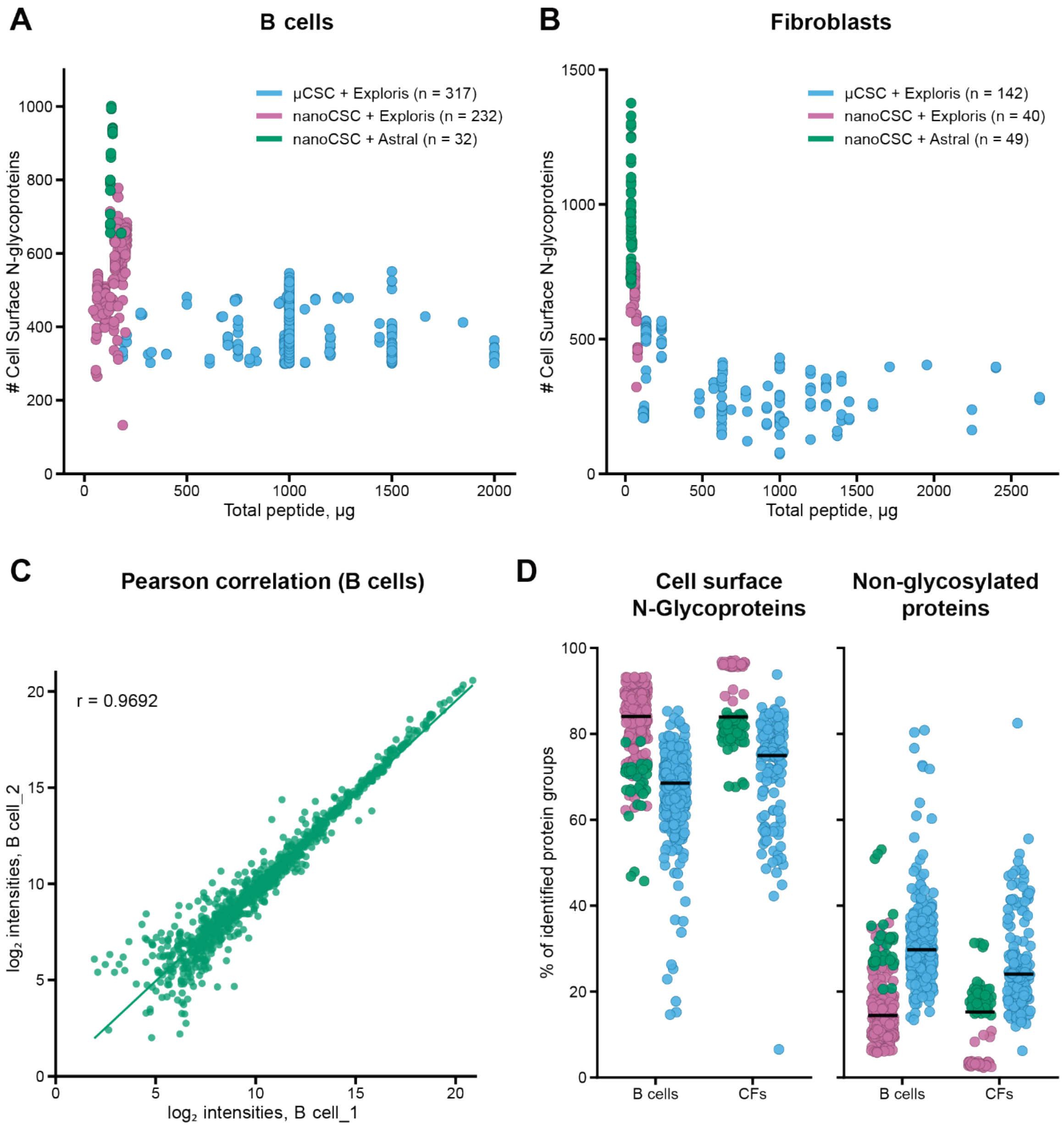


**Supplementary Figure 1. nanoCSC identifies more cell surface N-glycoproteins from lower starting material than μCSC, with high reproducibility.** A-B. Number of cell surface N-glycoproteins per run versus total peptide input in B cells (A) and CF (B). Each dot represents a run; colors identify the method and corresponding mass spectrometer used (μCSC + Exploris, blue; nanoCSC + Exploris, magenta-pink; nanoCSC + Astral (more sensitive MS), green). C. nanoCSC reproducibility in two separate analyses of B cells. $Log_2$ intensities of cell surface N-glycoproteins (Veneer High and Medium confidence) quantified in both B cell samples 1 and 2 (Pearson r = 0.9692; n = 843 protein groups). The line represents a linear fit. D. Percentage of identified protein types per run classified as cell surface N-glycoproteins (left) or non-glycosylated (right). Same colors and symbols as in A and B; black lines represent the median.

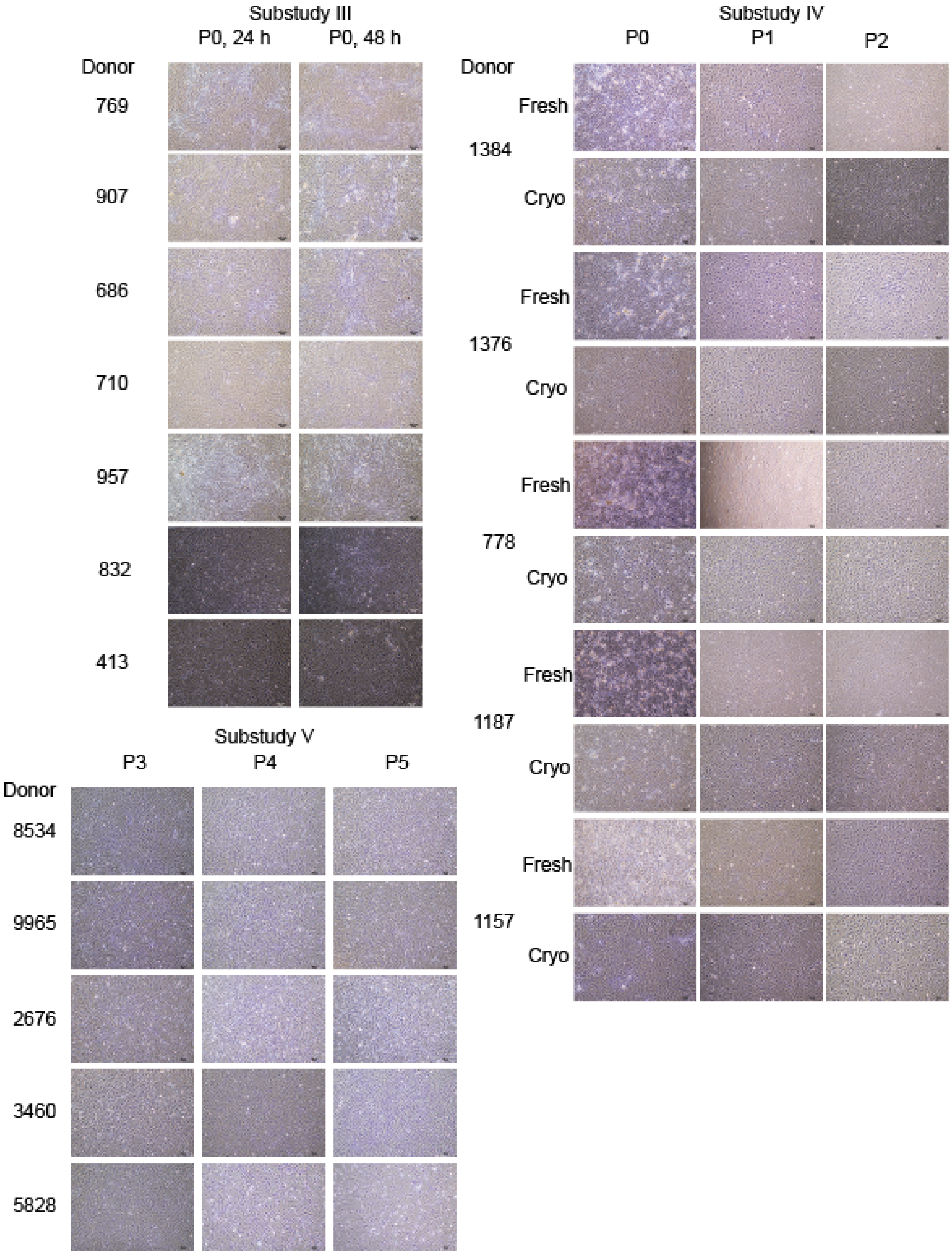

Substudy III
P0, 24 h
P0, 48 h
Donor
769
907
686
710
957
832
413
Substudy IV
P0
P1
P2
Donor
Fresh
1384
Cryo
Fresh
1376
Cryo
Fresh
778
Cryo
Fresh
1187
Cryo
Fresh
1157
Cryo
Substudy V
P3
P4
P5
Donor
8534
9965
2676
3460
5828

**Supplementary Figure 2. Brightfield images of cell cultures across the substudies.** Substudy III (top left): Fibroblast cultures from seven donors (769, 907, 686, 710, 957, 832, and 413) at passage 0, imaged 24 h and 48 h after plating. Substudy IV (right): Fibroblast cultures from five donors (1384,1376, 778, 1187, and 1157) established from fresh or cryopreserved (cryo) tissues, imaged at passages 0, 1, and 2. Substudy V (bottom left): commercially obtained Lonza fibroblast cell line from five donors (8534, 9965, 2676, 3460, and 5828) at passages 3, 4, and 5. Images were acquired at 40x magnification. Scale bar = 100 μm.

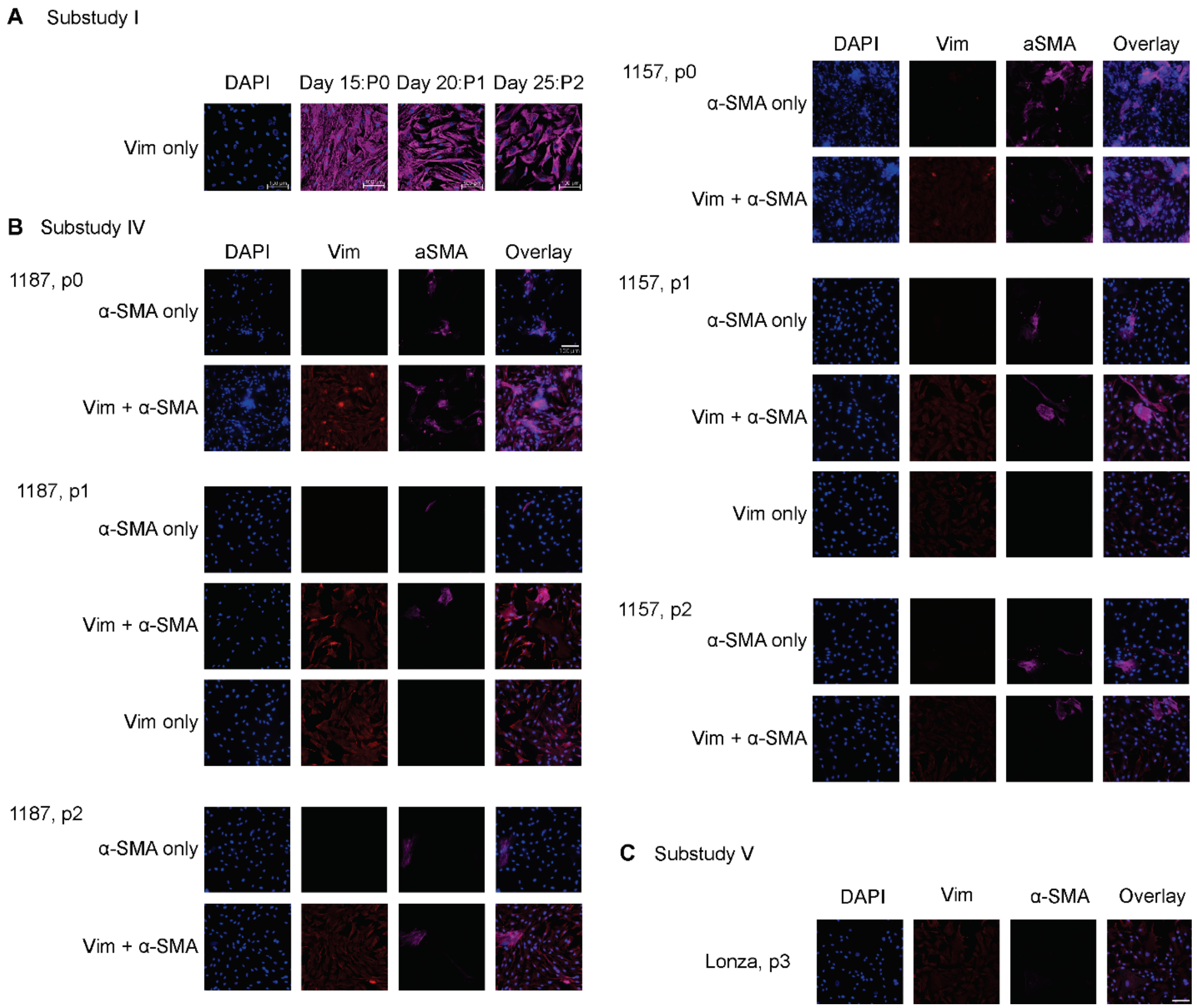


**Supplementary Figure 3. Immunofluorescent staining for vimentin (Vim) and α-smooth muscle actin (ACTA2, α-SMA) across passages.** (A) Substudy I: Fibroblasts from non-failing donor apex at passages 0, 1, and 2. Cells were stained with Vim and Hoechst counterstain. (B) Substudy IV: Fibroblasts from donors 1187 (left) and 1157 (right) at passages 0, 1, and 2. Cells were stained with α-SMA only, Vim and α-SMA, or Vim only. Each row includes DAPI nuclear counterstain, Vim, α-SMA, and merged overlay. (C) Substudy V: commercially obtained Lonza fibroblast cell line at passage 3 stained with DAPI, Vim, and α-SMA. Images were acquired at 40x magnification. Scale bar = 100 μm.

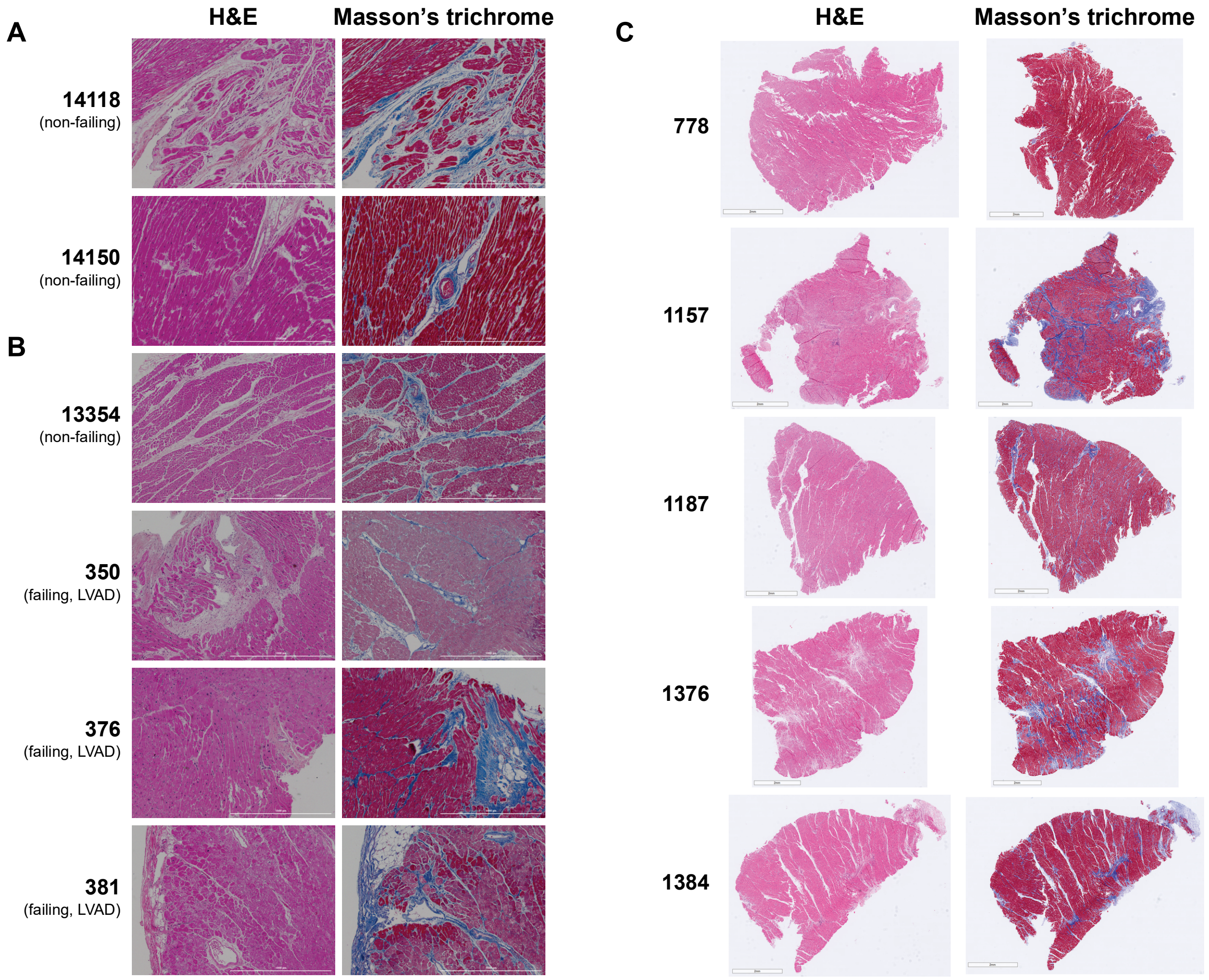


**Supplementary Figure 4. Assessment of cardiac fibrosis by histology.** Representative H&E (left) and Masson's trichrome (MTC, right) images; collagen deposition (blue on MTC) marks the fibrotic areas. A. Substudy I (non-failing donor apex): 14118 and 14150. B. Substudy II (failing vs. non-failing donor apex): non-failing (14118, 14150, 13354) and failing/LVAD (350, 376, 381); 14118 and 14150 (boxed) are shared with panel A. C. Substudy IV (whole-section left ventricle): 778, 1157, 1187, 1376, 1384; collagen fiber content was quantified from MTC images using CT-FIRE. Scale bars: apex (A, B), 1000 μm; left ventricle (C), 2 mm.

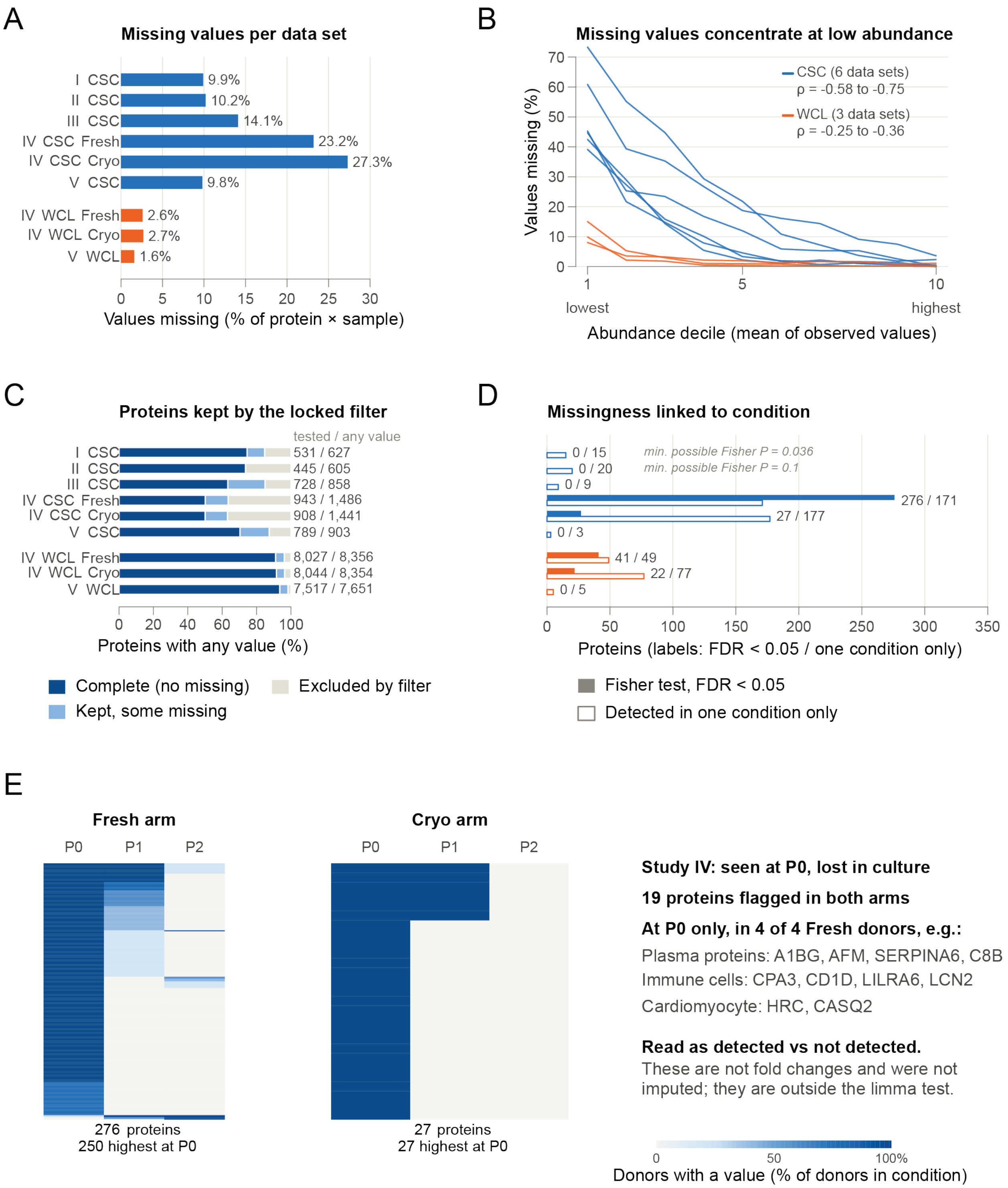


**Supplementary Figure 5. Missing values across the five studies.** A. Percentage of missing protein × sample values in each data set, before filtering (CSC 9.8–27.3%; whole cell lysate, WCL, 1.6–2.7%). CSC and WCL data sets are shown in blue and orange, respectively. B. Percentage of values missing in each tenth of proteins ranked by mean observed abundance; each line is one data set. ρ is the Spearman correlation between percent missing and mean abundance (CSC -0.58 to -0.75; WCL -0.25 to -0.36). C. Proteins observed in each data set that were complete in all samples, kept by the prespecified filter with some values missing, or excluded. The filter required quantification in a minimum number of donors in every condition (Study I ≥2, Substudy II ≥3, Substudy III ≥4, Substudy IV ≥3, Substudy V ≥3). Numbers on the right are proteins tested/proteins with any value. D. Proteins whose missingness depended on condition (Fisher exact test, Benjamini–Hochberg FDR < 0.05; filled bars) and proteins detected in only one condition (open bars). For Studies I and II, the smallest P value the group sizes allow is shown. E. For Substudy IV, the proteins with condition-linked missingness (Fresh arm 276, Cryo arm 27; 19 in both) are shown as the fraction of donors with a value at P0, P1, P2; rows are proteins sorted by detection pattern. 250 of 276 (Fresh) were most often detected at P0; 27 of 27 (Cryo) were most often detected at P0. No values were imputed.

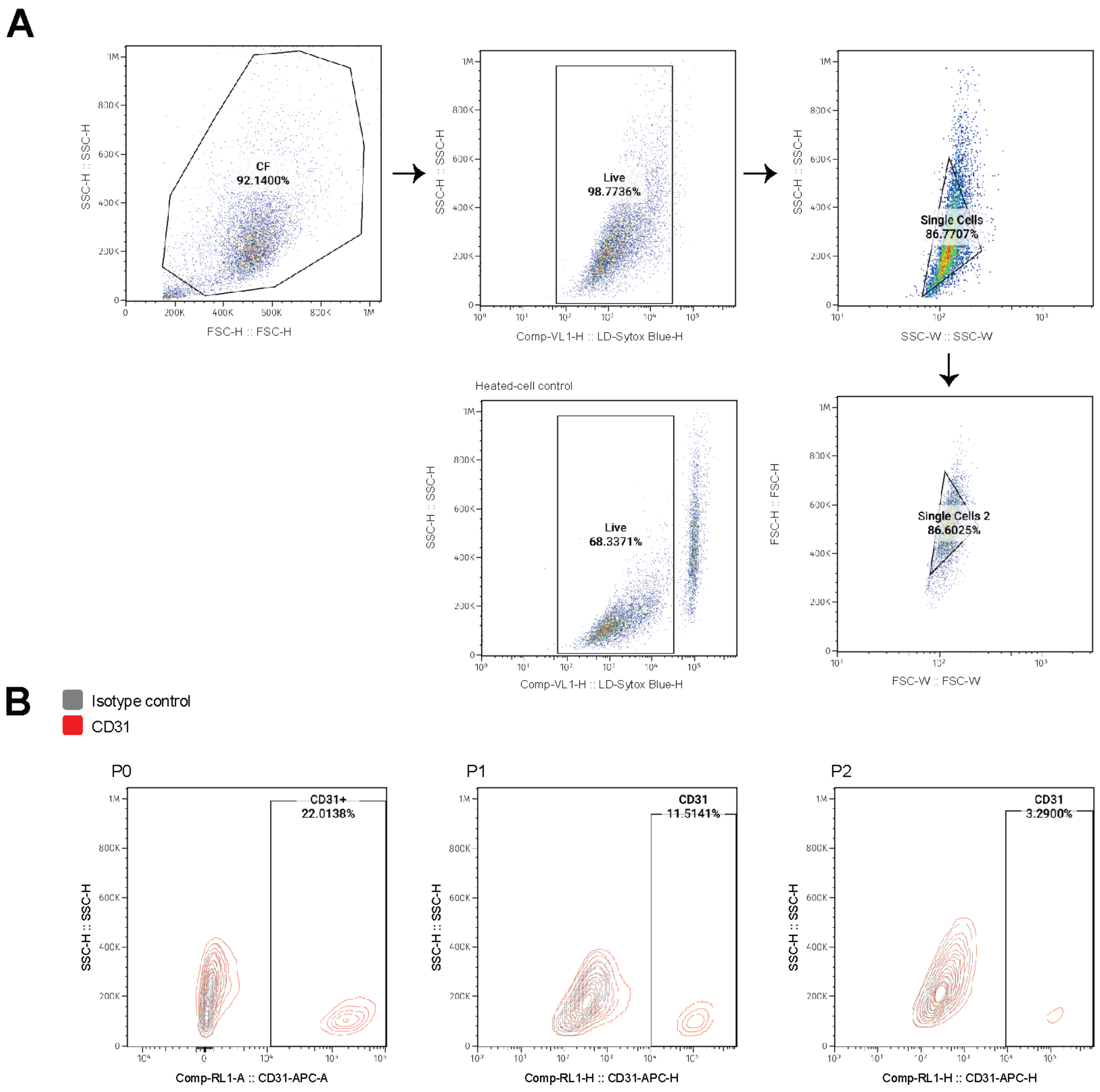


**Supplementary Figure 6. Flow cytometry of PECAM1 (CD31) in CF at P0, P1, and P2 (Substudy I).** A. Gating strategy. Cells were gated on forward and side scatter height (FSC-H vs. SSC-H) to exclude debris and on FSC-W vs. FSC-H to select single cells. PECAM1-positive cells were defined relative to the isotype control. B. Density plots of PECAM1 staining in CF at P0, P1, and P2 from one donor, with the percentage of PECAM1-positive live single cells shown in each gate. Antibody, fluorophore, and instrument details are provided in Supplementary Methods and Supplementary Table 2.

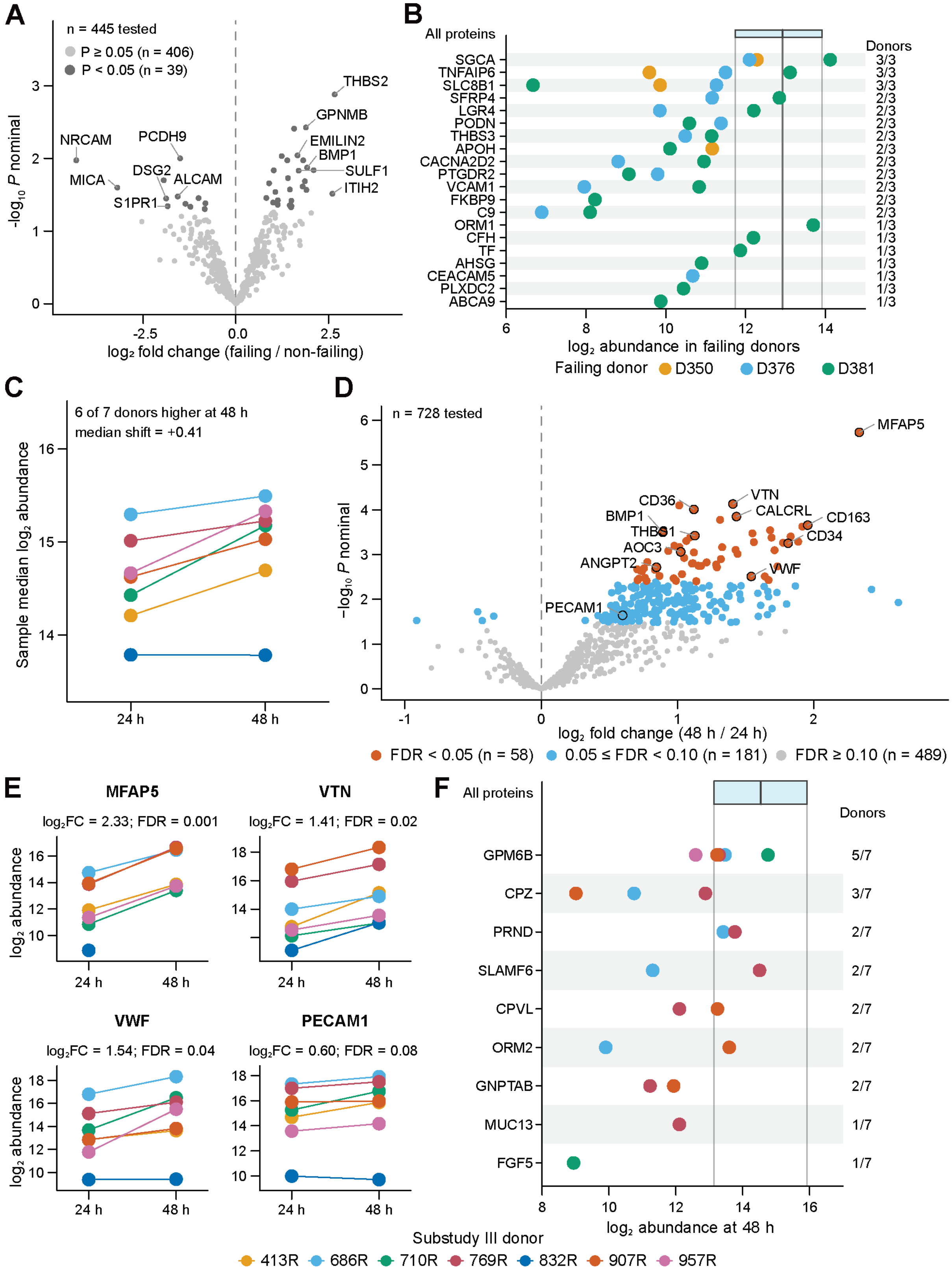


**Supplementary Figure 7. Surfaceome of CF from failing and non-failing hearts at P0 (Substudy II) and after 24 h or 48 h initial adhesion (Substudy III)**. A Volcano plot of $\log_2$

fold change (failing/non-failing) against -$\log_{10}$ nominal P value from limma for proteins tested in Substudy II; points are colored by nominal P value (dark gray, $P < 0.05$; light gray, $P \geq 0.05$). Selected proteins with $P < 0.05$ are labeled. B. Proteins detected in at least one failing donor and in no non-failing donor. Each symbol is the $\log_2$ abundance in one failing donor in which the protein was detected; filled symbols, detected in at least two of three failing donors; open symbols, one donor. The number of failing donors with detection is shown at right, and dotted lines separate proteins detected in three, two and one donor. The shaded band and vertical line mark the interquartile range and median abundance of all quantified proteins in Substudy II. C. Sample median $\log_2$ abundance of each Substudy III donor after 24 h and 48 h initial adhesion. D. Volcano plot of $\log_2$ fold change (48 h/24 h) against $-\log_{10}$ nominal P value from limma, colored by Benjamini–Hochberg FDR: vermillion, FDR < 0.05; blue, 0.05 ≤ FDR < 0.10; gray, FDR ≥ 0.10. Selected proteins are labeled. E. $\text{Log}_2$ abundance of MFAP5, VTN, VWF and PECAM1 in each donor at 24 h and 48 h, with the limma $\log_2$ fold change and FDR. F. Proteins detected at 48 h and in no donor at 24 h, shown as in B; filled symbols, detected in at least four of seven donors.

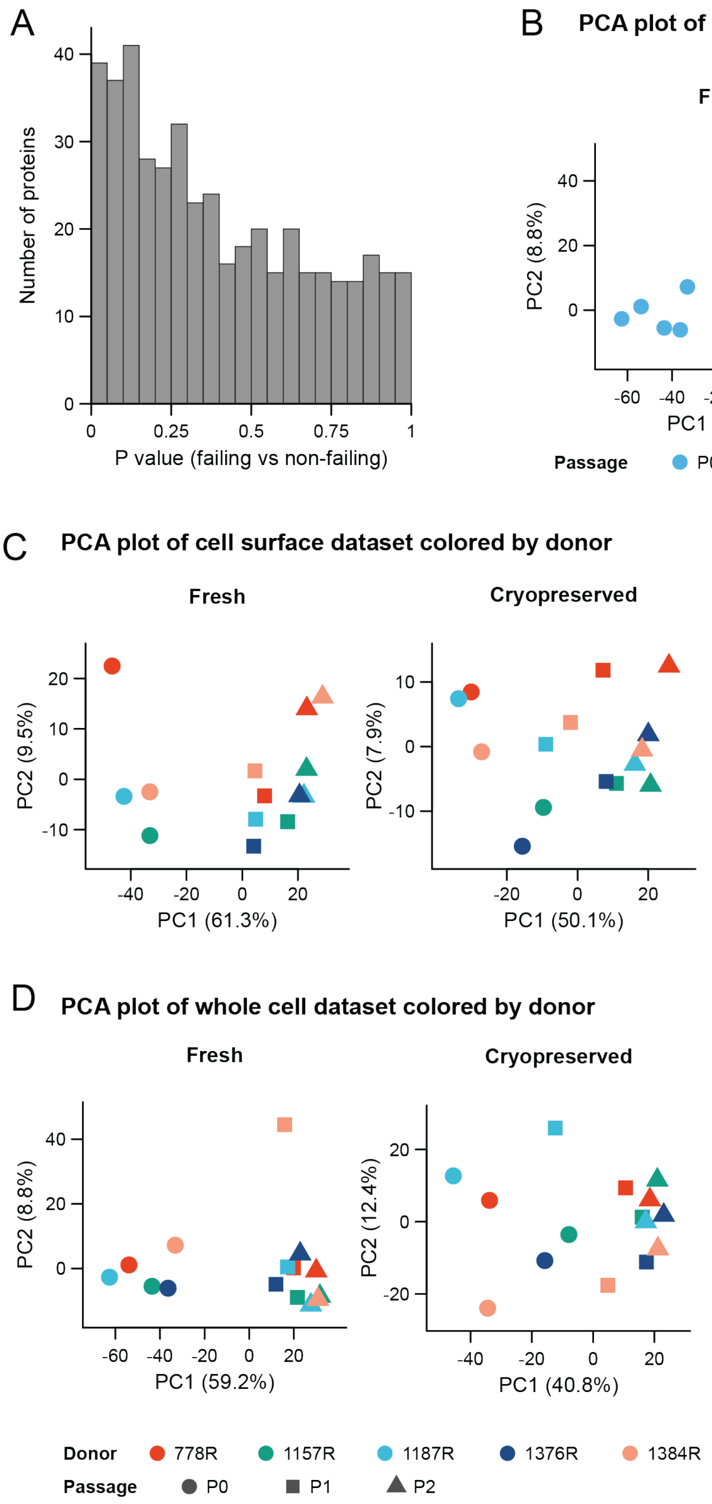


**Supplementary Figure 8. P-value distribution for Substudy II and principal component analysis of Substudy IV.** A. Distribution of P-values for failing vs. non-failing hearts in Substudy II (445 tested proteins). B-D. Principal component analysis of Substudy IV fresh and cryopreserved arms using proteins quantified in every sample: whole cell data colored by passage (B), cell surface data colored by donor (C), and whole cell data colored by donor (D).

Each point represents one sample; in C and D, symbols indicate passage (circle, P0; square, P1; triangle, P2). The percentage of variance explained by PC1 and PC2 is shown on the axes.

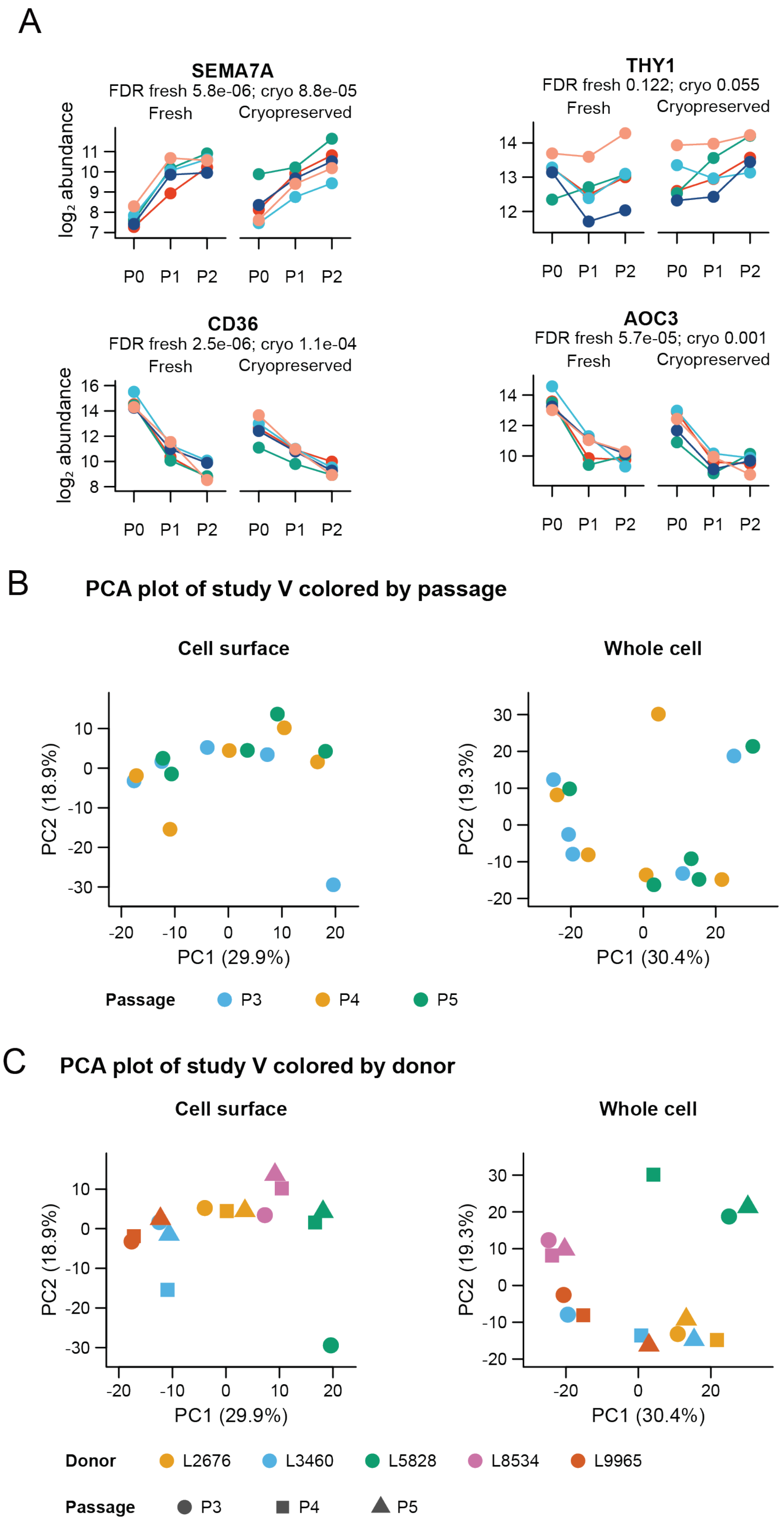
A
SEMA7A
FDR fresh 5.8e-06; cryo 8.8e-05
Fresh
Cryopreserved
log₂ abundance
P0 P1 P2
THY1
FDR fresh 0.122; cryo 0.055
CD36
FDR fresh 2.5e-06; cryo 1.1e-04
AOC3
FDR fresh 5.7e-05; cryo 0.001
B
PCA plot of study V colored by passage
Cell surface
Whole cell
PC2 (18.9%)
PC1 (29.9%)
PC2 (19.3%)
PC1 (30.4%)
Passage
P3
P4
P5
C
PCA plot of study V colored by donor
Donor
L2676
L3460
L5828
L8534
L9965

**Supplementary Figure 9. Whole cell trajectories in Substudy IV and principal component analysis of Substudy V.** A. $Log_2$ abundance of SEMA7A, THY1, CD36, and AOC3 in whole cell data at P0, P1, and P2 in each donor of the fresh and cryopreserved arms of Substudy IV. Each line is one donor**.** The FDR across passages in each arm is shown below each protein name. B-C. Principal component analysis of cell surface and whole cell protein abundance in Substudy V (commercially sourced CF, P3–P5), using proteins quantified in every sample. Each dot is one sample, colored by passage (B) or by donor, with symbols indicating passage (circle, P3; square, P4; triangle, P5) (C). The percentage of variance explained by PC1 and PC2 is shown on the axes.

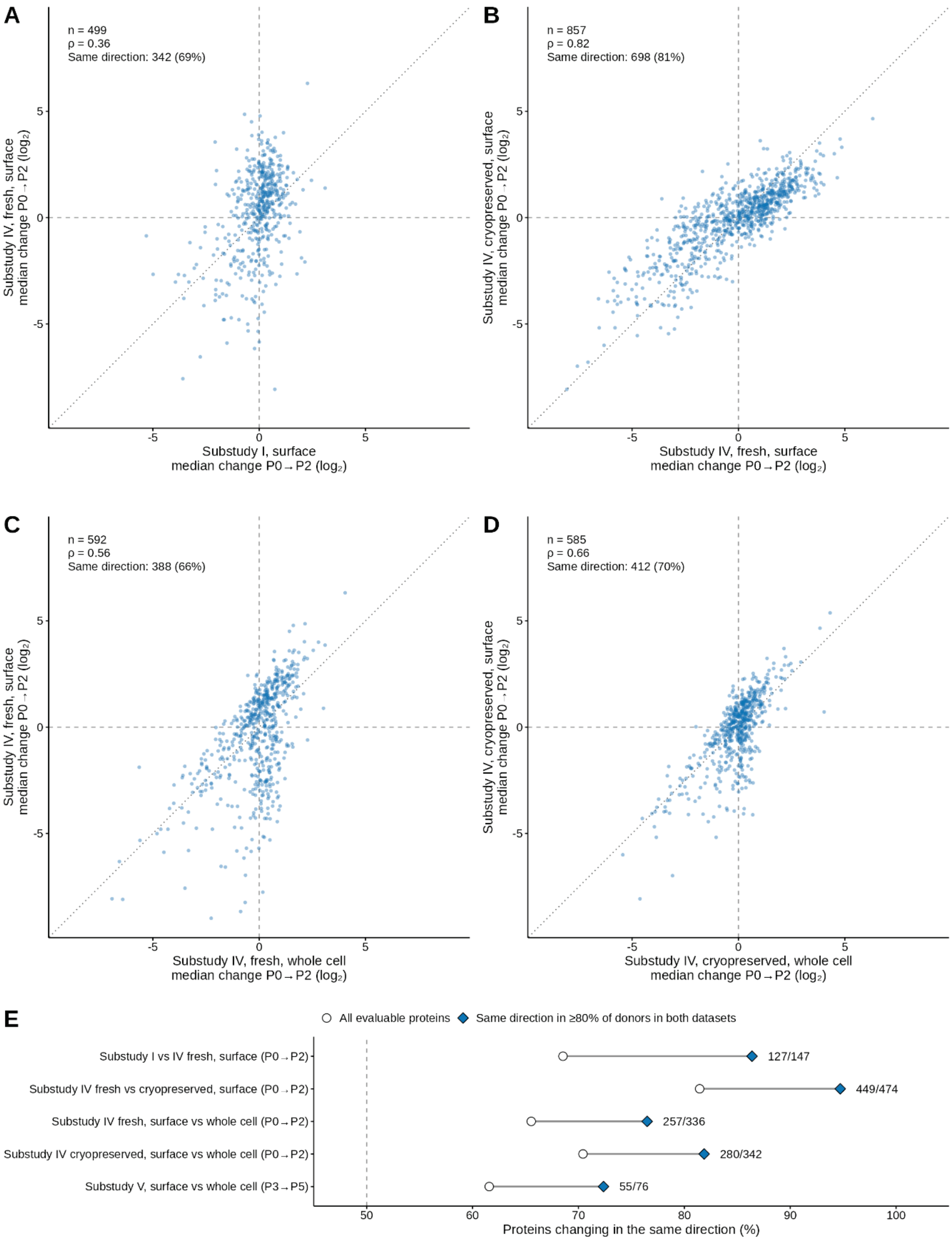
A
n = 499
ρ = 0.36
Same direction: 342 (69%)
Substudy IV, fresh, surface median change P0→P2 (log2)
Substudy I, surface median change P0→P2 (log2)
B
n = 857
ρ = 0.82
Same direction: 698 (81%)
Substudy IV, cryopreserved, surface median change P0→P2 (log2)
Substudy IV, fresh, surface median change P0→P2 (log2)
C
n = 592
ρ = 0.56
Same direction: 388 (66%)
Substudy IV, fresh, surface median change P0→P2 (log2)
Substudy IV, fresh, whole cell median change P0→P2 (log2)
D
n = 585
ρ = 0.66
Same direction: 412 (70%)
Substudy IV, cryopreserved, surface median change P0→P2 (log2)
Substudy IV, cryopreserved, whole cell median change P0→P2 (log2)
E
All evaluable proteins
Same direction in ≥80% of donors in both datasets
Substudy I vs IV fresh, surface (P0→P2)
Substudy IV fresh vs cryopreserved, surface (P0→P2)
Substudy IV fresh, surface vs whole cell (P0→P2)
Substudy IV cryopreserved, surface vs whole cell (P0→P2)
Substudy V, surface vs whole cell (P3→P5)
127/147
449/474
257/336
280/342
55/76
Proteins changing in the same direction (%)

**Supplementary Figure 10. Agreement of passage-associated changes across cohorts, tissue preparations, and assays.** A-D. Each point represents one protein. The axes show the median within-donor change in $\log_2$ abundance from P0 to P2, for proteins measured at both passages in at least 60% of donors in both datasets. Dashed lines mark zero change and the dotted line marks identity. Each panel shows the number of proteins (n), Spearman correlation (ρ), and number and percentage of proteins that changed in the same direction. A. Cell surface data from Substudy I and the fresh arm of Substudy IV. B. Cell surface data from the fresh and cryopreserved arms of Substudy IV. C-D. Whole cell and cell surface data from the fresh (C) and cryopreserved (D) arms of Substudy IV. (E) Percentage of proteins that changed in the same direction in each comparison, for all evaluable proteins (open circles) and for proteins that changed in the same direction in at least 80% of donors in both datasets (filled diamonds). Numbers indicate the proteins in the second group that agreed in direction, out of the total. The last row compares surface and whole cell data in Substudy V from P3 to P5. The dashed line marks 50%.

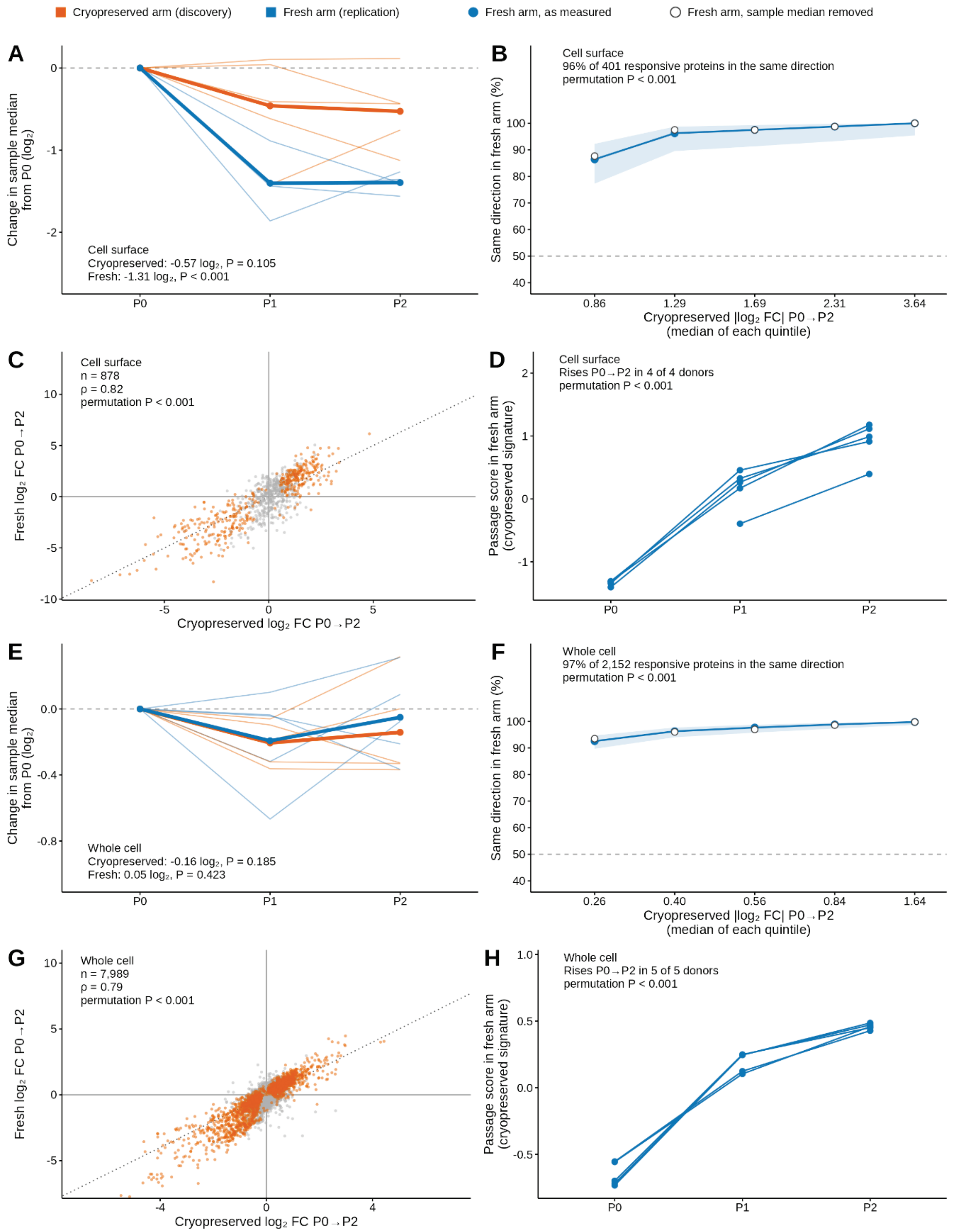


**Supplementary Figure 11. Agreement of passage-associated changes between CF from cryopreserved and fresh myocardium (Substudy IV).** Cell surface data are shown in A-D and whole cell data in E-H. The cryopreserved arm was used to define passage-responsive proteins (FDR < 0.05, P0 vs P2), and the fresh arm was used to test them. A, E. Change in sample median $\log_2$ abundance from P0, calculated before normalization, for each donor (thin lines) and the arm mean (thick lines) in the cryopreserved (orange) and fresh (blue) arms. The values are the mean change from P0 to P2, with P values from a test of passage with donor as a blocking factor. B, F. Percentage of responsive proteins that changed in the same direction in the fresh arm, in quintiles of the absolute $\log_2$ fold change in the cryopreserved arm. Filled circles show the data as measured, with 95% confidence intervals (shaded); open circles show the data after each sample's median was removed. The dashed line marks 50%, the agreement expected by chance. C, G. Limma $\log_2$ fold change from P0 to P2 in the cryopreserved and fresh arms for all proteins tested in both. Responsive proteins are shown in orange and the dotted line marks identity. D, H. Passage score for each fresh-arm sample. The score is the mean within-donor-centered abundance of the responsive proteins, each signed by its direction of change in the cryopreserved arm. Lines connect samples from the same donor; one donor had no P0 cell surface sample. Permutation P values come from 1,000 shuffles of passage labels within donors in the fresh arm.

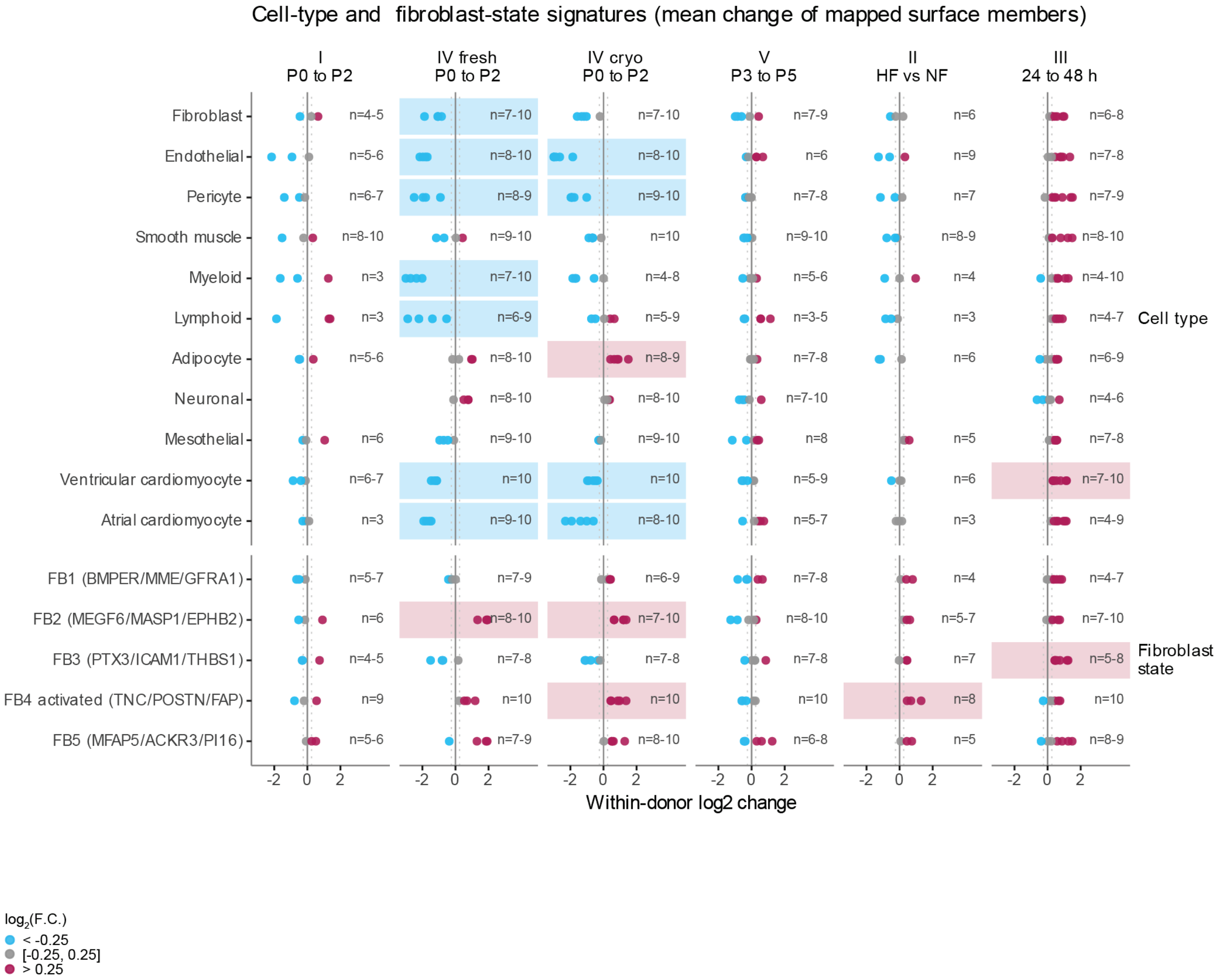


**Supplementary Figure 12.** Mean within-donor $\log_2$ change of the members of each cell-type and fibroblast-state signature quantified at both endpoints (at least three), shown as in Figure 5; n, members contributing per donor. FB6 and FB7 are not shown because their top markers duplicate endothelial and cardiomyocyte signatures.

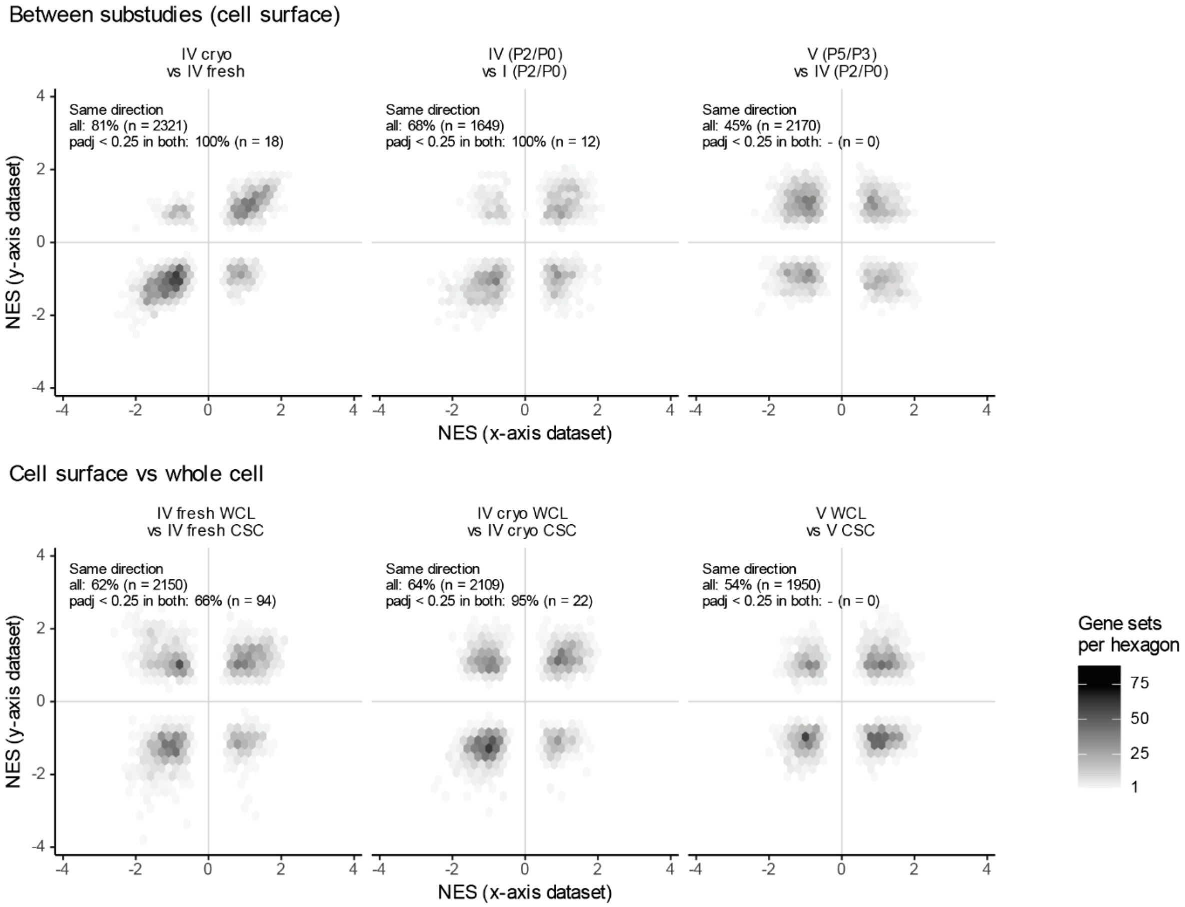


**Supplementary Figure 13.** Gene sets binned by their NES in two datasets: between substudies at the cell surface (top), cell surface versus whole cell data (middle), and the same with whole cell data restricted to proteins tested at the surface (bottom). Labels indicate the percentage of gene sets with NES of the same sign for all sets and for sets with adjusted P < 0.25 in both datasets.

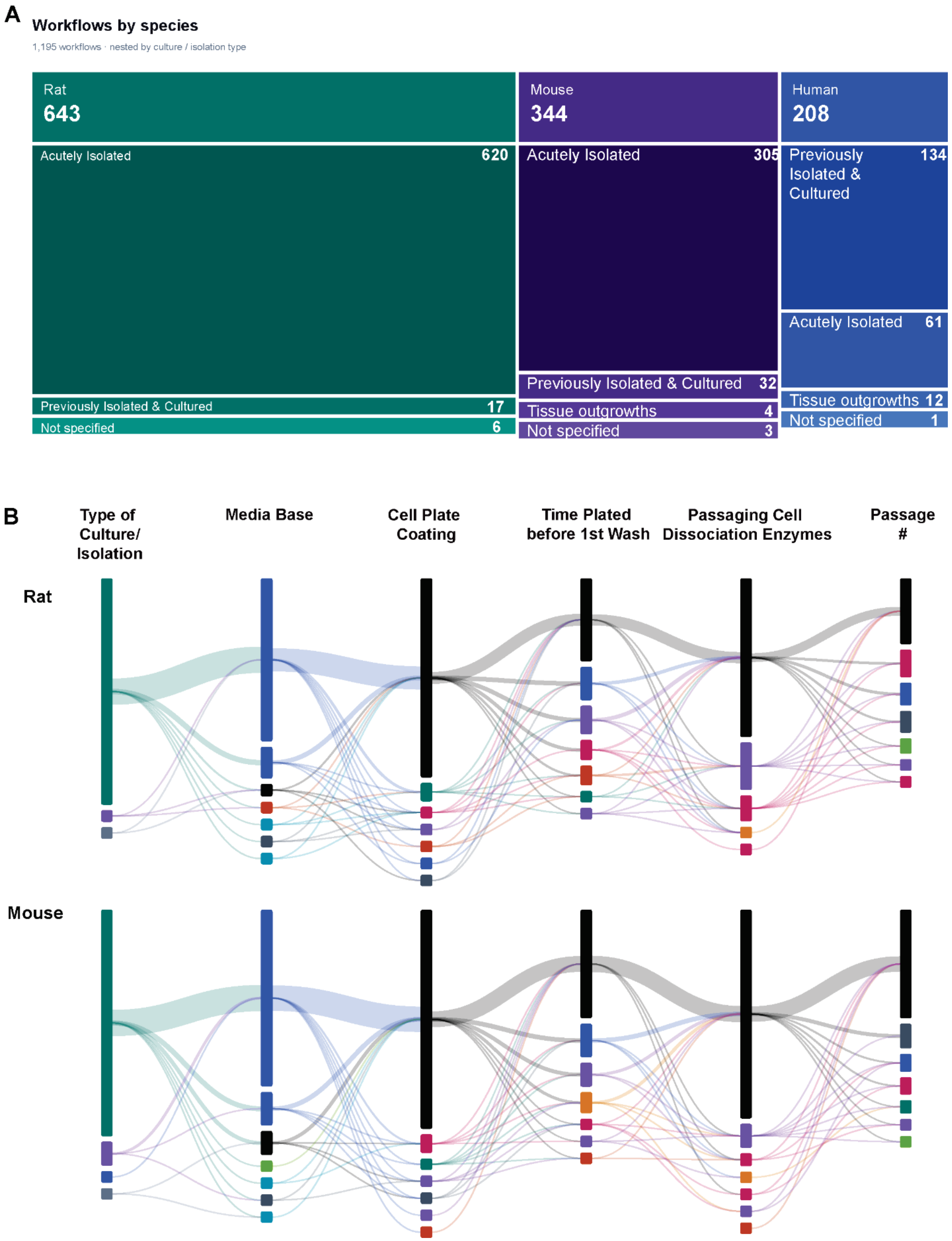


**Supplementary Figure 14. CF culture practices reported across 30 years of literature for mouse and rat studies.** A. A nested view showing how extracted culture workflows divide by species and culture or isolation approach. B. Sankey diagram of the combinations of six culture variables (cell source, basal medium, plate coating, initial adhesion time, dissociation enzymes, and passage) among studies using rat or mouse CF.
